# Comprehensive Interactome Mapping of the DNA Repair Scaffold SLX4 using Proximity Labeling and Affinity Purification

**DOI:** 10.1101/2022.09.19.508447

**Authors:** Camila M. Aprosoff, Boris J.A. Dyakov, Vivian H.W. Cheung, Cassandra J. Wong, Mikaela Palandra, Anne-Claude Gingras, Haley D.M. Wyatt

**Affiliations:** Department of Biochemistry, University of Toronto, Toronto, ON, M56 1A8, Canada; Department of Molecular Genetics, University of Toronto, Toronto, ON, M56 1A8, Canada; Lunenfeld-Tanenbaum Research Institute, Mount Sinai Hospital, Sinai Health, 600 University Avenue, Toronto, M5G 1X5, Canada; Canada Research Chairs Program, Temerty Faculty of Medicine, University of Toronto, Toronto, ON, M5S 1A8, Canada

**Keywords:** SLX4, DNA repair, genome stability, proteomics, BioID, AP-MS

## Abstract

The DNA repair scaffold SLX4 has pivotal roles in cellular processes that maintain genome stability, most notably homologous recombination. Germline mutations in *SLX4* are associated with Fanconi anemia, a disease characterized by chromosome instability and cancer susceptibility. The role of mammalian SLX4 in homologous recombination depends critically on binding and activating structure-selective endonucleases, namely SLX1, MUS81-EME1, and XPF-ERCC1. Increasing evidence indicates that cells rely on distinct SLX4-dependent complexes to remove DNA lesions in specific regions of the genome. Despite our understanding of SLX4 as a scaffold for DNA repair proteins, a detailed repertoire of SLX4 interactors has never been reported. Here, we provide the first comprehensive map of the human SLX4 interactome using proximity-dependent biotin identification (BioID) and affinity purification coupled to mass spectrometry (AP-MS). We identified 237 high-confidence interactors, of which the vast majority represent novel SLX4 binding proteins. Network analysis of these hits revealed pathways with known involvement of SLX4, such as DNA repair, and novel or emerging pathways of interest, including RNA metabolism and chromatin remodeling. In summary, the comprehensive SLX4 interactome we report here provides a deeper understanding of how SLX4 functions in DNA repair while revealing new cellular processes that may involve SLX4.

## INTRODUCTION

The accurate duplication and faithful transmission of genetic information into progeny is vital for cell growth and survival. These processes are threatened by DNA damage, which frequently results from exposure to factors in the environment (e.g., ultraviolet radiation) or from within cells (e.g., aldehydes produced by lipid peroxidation and alcohol metabolism) [1]. One of the most dangerous types of damage is a DNA double-strand break (DSB), which can result from external sources such as ionizing radiation. However, DSBs also arise during normal biological processes, including V(D)J recombination and DNA replication, the latter occurring when the replisome encounters an unrepaired single-stranded break or after nucleolytic processing of dysfunctional replication forks. Importantly, unrepaired DNA damage is a major driving force for genome instability and cancer development [2].

Cells use sophisticated DNA repair networks to counteract the harmful effects of genotoxic agents, thus safeguarding genome integrity. The two major repair pathways for repairing DSBs in eukaryotes are non-homologous end joining (NHEJ) and homologous recombination (HR). In its simplest form, NHEJ is a rapid process that entails DNA end modification and subsequent ligation [3]. This reaction depends minimally on sequence complementarity, meaning that NHEJ can occur throughout the cell cycle [4]. By contrast, HR requires extensive sequence homology between the broken DNA and a donor (or template) DNA molecule [5]. Homologous recombination is restricted to S and G2-phases of the cell cycle, with the sister chromatid generally used as the template for repair in somatic cells [4, 6]. The physiological importance of accurate DNA repair is underscored by the fact that germline mutations in DSB repair genes cause genome instability in numerous hereditary diseases associated with cancer predisposition and neurological defects [1].

Homologous recombination is an intricate and multistep process (for reviews, see [5, 7]). One of the first steps is DNA end resection, which generates extended 3’ single-stranded DNA (ssDNA) tails that are rapidly coated with the heterotrimeric RPA complex. Next, recombination ‘mediators’ displace RPA and promote the loading of RAD51 to form a dynamic nucleoprotein filament that invades duplex DNA to mediate homology search, resulting in a recombination intermediate called the displacement loop. If sufficient base-pairing occurs, the 3’-end of the invading strand engages a DNA polymerase, which extends the nascent strand using the donor DNA as a template. Although several downstream pathways exist, the classical DSB repair model entails the formation of four-stranded intermediates called Holliday junctions, which must be removed to complete HR (for reviews, see [5, 8]). Importantly, cells contain two distinct mechanisms to process Holliday junctions. The ‘dissolution’ pathway involves the BLM helicase in complex with DNA topoisomerase 3α (TOP3α), RMI1 and RMI2, and gives rise to non-crossover products. Alternatively, structure-selective endonucleases promote Holliday junction ‘resolution’ by introducing a pair of nicks across the helical axis, generating either crossover or non-crossover products [8]. Holliday junction resolvases in mammalian cells include GEN1 and a macromolecular nuclease complex built on a scaffold protein called SLX4 [9–11].

The role of mammalian SLX4 in HR depends critically on binding and activating structure-selective endonucleases, namely SLX1, MUS81-EME1, and XPF-ERCC1 [10–15]. These enzymes remove branched DNA structures that form during HR, such as Holliday junctions [16, 17]. This is important because persistent branched DNA structures impede accurate HR and chromosome segregation, leading to genome instability. Early clues about the scaffold function of mammalian SLX4 came from bioinformatics and affinity purification coupled to mass spectrometry (AP-MS) experiments, which revealed that SLX1, MUS81-EME1 and XPF-ERCC1 bind distinct regions of SLX4 (Figure 1A) [12–14]. Subsequent biochemical studies revealed that SLX4 activates and regulates its endonuclease partners. For example, SLX1 is catalytically inactive until it binds the conserved C-terminal domain (CCD) domain of SLX4, after which it is competent to cleave many types of branched DNA structures [10, 12–14]. Similarly, SLX4 stimulates XPF-ERCC1 to cleave replication forks that stall at protein-DNA adducts or interstrand crosslinks [18, 19]. When cells enter mitosis, phosphorylation of the SLX4 scaffold triggers the recruitment of MUS81-EME1, leading to the formation of a tri-nuclease complex called SMX (for <u>S</u>LX1-SLX4, <u>M</u>US81-EME1, and <u>X</u>PF-ERCC1) [10, 15]. Notably, SLX4 binding to MUS81-EME1 stimulates the cleavage of branched DNA structures that represent replication and recombination intermediates [15]. The current model is that SMX provides cells with a multifunctional nuclease that removes joint molecules prior to cytokinesis. The cell cycle-regulated assembly of SMX in mitosis is thought to prevent catastrophic cleavage of replicating DNA in S-phase [20–23]. As such, there is great interest in understanding the molecular mechanisms that regulate SMX (dis)assembly and activity.

**Figure 1.**
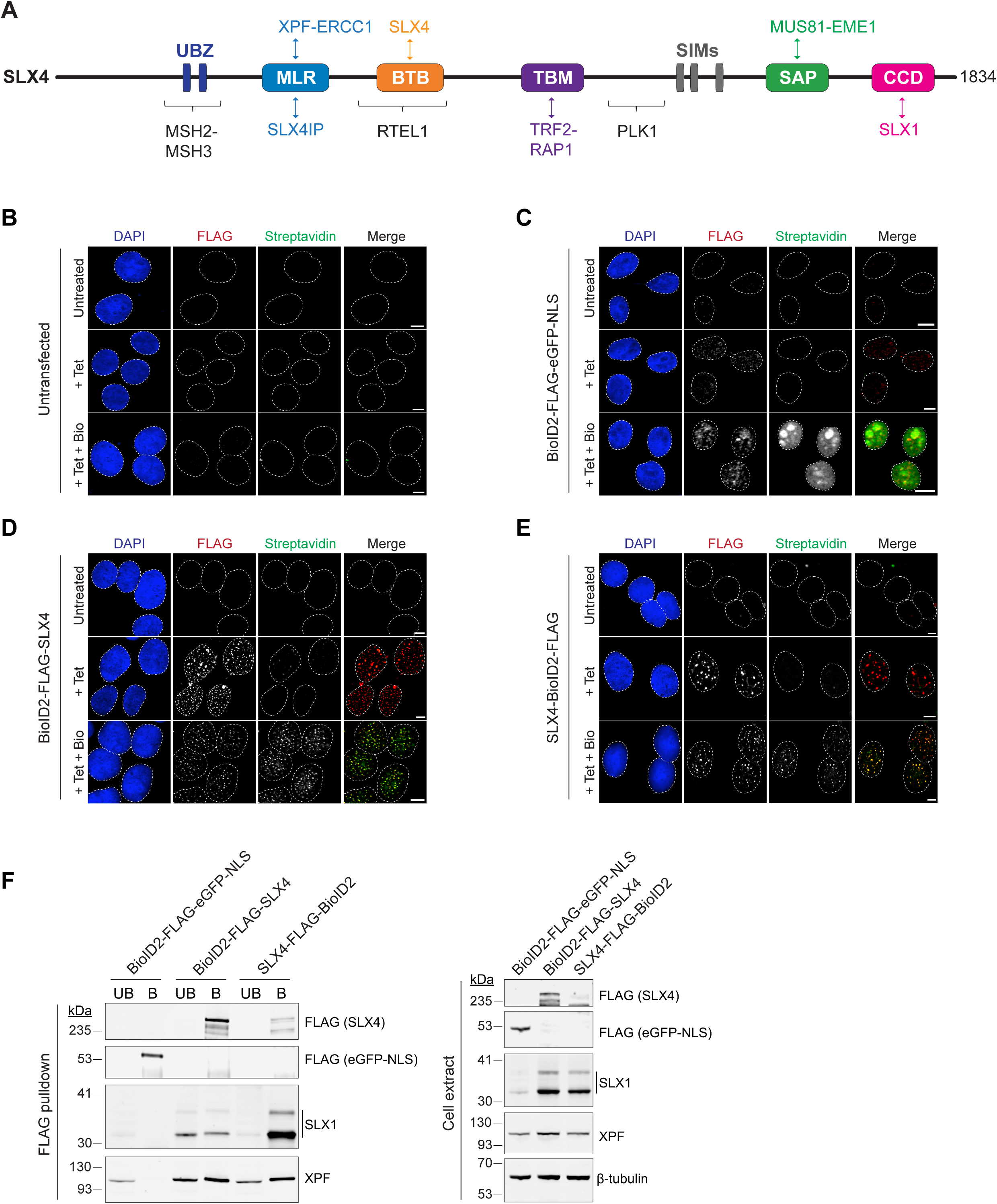
Nuclear localization and functionality of BioID2-FLAG-SLX4 and SLX4-FLAG-BioID2 in human cells. **(A)** Domain architecture of the human SLX4 scaffold highlighting known interaction partners and their binding regions. Abbreviations for protein domains: UBZ4, ubiquitin-binding zinc finger type 4; MLR, MUS312-MEI9 interaction-like region; BTB, Broad-Complex, Tramtrack and Bric a Brac; TBM, TRF2-binding motif; SIM, SUMO-interacting motif; SAP, SAF-A/B, Acinus and PIAS; CCD, conserved C-terminal domain. **(B-E)** Confocal immunofluorescence microscopy analysis of untransfected Flp-In T-Rex HEK293 cells **(B)**, or cells stably expressing BioID2-FLAG-eGFP-NLS **(C)**, BioID2-FLAG-SLX4 **(D)**, or SLX4-FLAG-BioID2 **(E)**. Cultures were left untreated (*top panels*) or incubated with tetracycline for 25 hr to induce construct expression (*middle panels*) and biotin for 8 hr to induce the biotinylation of proximal proteins (*bottom panels*). Cells were fixed and processed for immunofluorescence using anti-FLAG (red) and streptavidin-conjugated Alexa488 (green) antibodies. Dashed lines demark nuclear boundaries, as determined using DAPI staining. Scale bar represents 50 μM. **(F)** Western blot analysis of Flp-In T-Rex HEK293 cells treated with tetracycline for 25 hr to induce expression of the indicated construct. Cells containing BioID2-FLAG-eGFP-NLS were induced with 0.001 μg/mL tetracycline, whereas BioID2-FLAG-SLX4 and SLX4-FLAG-BioID2 were induced with 1 μg/mL tetracycline. FLAG-tagged proteins were immunoprecipitated from 2.5 mg of extract using α-FLAG beads and analyzed by western blotting for the indicated proteins (left). In parallel, 20 μg cell extract was analyzed by western blotting for the indicated proteins (right). β-tubulin was used as a loading control. Abbreviations: UB = unbound fraction, B = bound fraction on α-FLAG beads.

In addition to structure-selective endonucleases, SLX4 interacts with other proteins that have important roles in DNA repair and genome stability maintenance, such as SLX4IP, the core telomere-binding proteins TRF2 and TERF2IP (RAP1), and the mismatch repair complex MSH2-MSH3 (Figure 1A). In general, these interactions promote nucleolytic cleavage of branched DNA structures in different regions of the genome [17]. For example, through the interaction with TRF2, SLX4 negatively regulates telomere lengths in cells that use the alternative lengthening of telomeres (ALT) mechanism of telomere elongation [24, 25]. This mechanism involves SLX1-dependent cleavage of branched DNA structures that arise during telomere replication and recombination [24, 26]. SLX4IP is also required for telomere length maintenance in ALT cells, although the precise role remains controversial [27–29]. Recent work showed that the mismatch repair heterodimer MSH2-MSH3 targets SLX4 and its associated nucleases to cleave expanded trinucleotide repeats, which are linked to numerous neurodegenerative disorders including myotonic dystrophy and Huntington’s disease [30]. Conversely, the interaction between SLX4 and MSH2 inhibits the activity of MSH2-MSH6, which recognizes single mismatches and small insertions and deletions (indels) during mismatch repair [31]. This is the first example where SLX4 inhibits DNA repair, highlighting the gaps in our knowledge about the composition and functions of SLX4-dependent protein complexes.

Post-translational modifications (PTMs) have central roles in the initial recognition of DNA damage and the initiation and execution of DNA repair [32]. In general, PTMs facilitate rapid and reversible protein recruitment to, and extraction from, sites of DNA damage. The binding partners, subcellular localization, and recruitment of SLX4 to different types of DNA damage are regulated by PTMs including phosphorylation, ubiquitylation, SUMOylation, and PARylation [20, 33–36]. These studies illustrate the complex and multilayer regulatory networks that direct SLX4 complexes to distinct functional contexts.

Despite the well-established role of SLX4 in HR and genome stability maintenance, the full repertoire of SLX4 binding proteins has not been investigated. Previous high-throughput studies relied exclusively on AP-MS to identify SLX4 interactors [13, 14, 33, 37]. Although these studies paved the way for our understanding of SLX4-containing complexes, one limitation of AP-MS (like most methods that rely on biochemical isolation of intact protein complexes) is its poor ability to capture weak, transient, or low-abundance interactions [38]. This technique is also susceptible to false positives caused by perturbation of cell compartments during lysis, allowing for interactions between proteins normally sequestered in different organelles [39].

In recent years, proximity-dependent labeling has emerged as a powerful method to gain deeper insight into protein networks. Proximity-dependent biotin identification (BioID) is method for identifying proximal protein-protein associations in the context of living cells (Figure S1A) [40, 41]. The protein of interest (bait) is fused to an abortive biotin ligase, resulting in the covalent biotinylation of lysine residues in neighboring proteins (prey) within nanometers of the bait [42–44]. Biotinylated proteins are captured on streptavidin-conjugated beads, alleviating the need for an intact bait-prey complex during cell lysis and bait isolation (Figure S1A). The main advantage of this technique is its ability to capture both direct and indirect interactors of the bait, as well as other proteins found in the vicinity of the tagged protein [40, 41]. Earlier studies employing BioID relied on the *Escherichia coli* BirA* biotin ligase tag (an enzyme with a single mutation at R118G), but newer enzymes have since been adapted that may prove useful in certain contexts [45]. The BioID2 tag uses the smaller *Aquifex aeolicus* biotin ligase, mutated to include an R40G substitution in the catalytic domain for abortive biotinylation [43]. Since *A. aeolicus* biotin ligase naturally lacks the DNA-binding domain, we reasoned it would be more appropriate for obtaining a good signal-to-noise ratio with nuclear and chromatin-associated proteins. Proximity-dependent labeling has become increasingly popular for detecting weak or transient interactions [46–48] and for determining the proteomes of cellular structures or compartments, such as the nuclear envelope [42], the nuclear pore complex [44], and cytosolic membraneless organelles [49]. Additionally, Go et al. created a human cell map using proximity-dependent biotinylation to profile 192 protein markers from 32 different subcellular compartments, including nuclear bodies, nucleoli, and chromatin [50].

Given that SLX4 is a scaffold for DNA repair and chromatin-associated proteins, we reasoned that it would be valuable to investigate the SLX4 interactome using both proximity-dependent biotin identification (BioID, using BioID2) and AP-MS [38, 39, 51]. Here, we present the first comprehensive map of the SLX4 interactome, confirming known partners and revealing a plethora of novel and unexpected interactors. By performing gene ontology (GO) and clustering analyses, we identified potential roles for SLX4 in biological functions that have not been previously ascribed to the scaffold. We identified several candidate interactors that could regulate SLX4 PTMs, including phosphorylation, ubiquitylation and SUMOylation. We also observed a significant number of high-confidence interactors involved in chromatin remodeling and transcription, suggesting that these could represent understudied functions of SLX4. Overall, our study provides a rich resource of SLX4 interactors in undamaged human cells and a starting point for future studies aimed at understanding connections between SLX4 and processes beyond DNA repair.

## EXPERIMENTAL PROCEDURES

### Plasmids

Gateway entry constructs encoding *SLX4* (accession number 84464) were generated using Gateway BP Clonase II, as per the manufacturer’s instructions (Invitrogen, ThermoFisher Scientific). For N-terminal tagging, the start codon was mutated to alanine using the Q5 Site-Directed Mutagenesis Kit (NEB). For C-terminal tagging, the stop codon was deleted using the Q5 Site-Directed Mutagenesis Kit (NEB). Plasmids were transformed into *E. coli* One Shot ccdB Survival 2 TI^R^ competent cells (Invitrogen, ThermoFisher Scientific) and selected on LB agar supplemented with 50 µg/mL kanamycin. 3×FLAG-BioID2 constructs were generated by performing a Gateway LR Clonase II reaction (Invitrogen, ThermoFisher Scientific) into the pDEST-pcDNA5-FRT/TO-BioID2-3×FLAG backbone (Hesketh et al., 2017) (with fusion of the marker at either the N or C terminus), as per the manufacturer’s instructions. Plasmids were transformed into *E. coli* XL10 Gold (Agilent Technologies) and colonies were selected on LB agar supplemented with 100 µg/mL ampicillin. Constructs were validated by Sanger sequencing (ACGT Corp., Toronto).

Negative control plasmids included the following: pDEST-pcDNA5-FRT/TO-BioID2-3×FLAG (BioID2-FLAG), pDEST-pcDNA5-FRT/TO-BioID2-3×FLAG-eGFP (BioID2-FLAG-eGFP), and pDEST-pcDNA5-FRT/TO-BioID2-3×FLAG-eGFP-NLS (BioID2-FLAG-eGFP-NLS), where eGFP is enhanced green fluorescent protein and NLS is the SV40-nuclear localization sequence [38].

### Human cell lines and culture conditions

All Flp-In T-REx human embryonic kidney 293 (HEK293) cells were cultured in high-glucose Dulbecco’s Modified Eagle’s Medium (DMEM) containing L-glutamine and 110 mg/L sodium pyruvate (Gibco, ThermoFisher Scientific) supplemented with 10% tetracycline-free fetal bovine serum (FBS) (Wisent, Inc.), 100 U/mL penicillin and 100 μg/mL streptomycin (Gibco, ThermoFisher Scientific). Untransfected Flp-In T-REx HEK293 cells were cultured under selection with 4 µg/mL blasticidin (Gibco, ThermoFisher Scientific) and 50 µg/mL zeocin (Invitrogen, ThermoFisher Scientific). The growth media for stably transfected cell cultures contained 4 μg/mL blasticidin (Gibco, ThermoFisher Scientific) and 100 µg/mL hygromycin (Invitrogen, ThermoFisher Scientific). Two 15-cm plates of cells were used for each BioID2 and AP-MS experimental replicate. All cultures were maintained at 37°C in a humidified chamber containing 6% CO_2_.

### Preparation of biotin-depleted FBS

To prepare biotin-depleted FBS, 50 mL of tetracycline-free FBS (Wisent, Inc.) was incubated with 1 mL of Streptavidin Sepharose HP (Cytiva) overnight at 4°C and then filtered through a 0.22 μm polyethersulfone (PES) filter (Nalgene, ThermoFisher Scientific).

### Generation of stable cell lines

To generate stable cell lines, Flp-In T-REx human embryonic kidney 293 (HEK293) cells were grown in 6-cm plates and co-transfected with plasmids pDEST-pcDNA5-FRT/TO, encoding a specific BioID2 construct, and pOG44 Flp-Recombinase (at 1:9 w/w ratio) using Lipofectamine 3000 (Invitrogen, ThermoFisher Scientific), according to the manufacturer’s instructions. At 24 hr post-transfection, the growth media was replaced with high-glucose Dulbecco’s modified Eagle’s medium containing L-glutamine and 110 mg/L sodium pyruvate (Gibco, ThermoFisher Scientific) and 10% tetracycline-free fetal bovine serum (Wisent, Inc.). At 48 hr post-transfection, cells were trypsinized and seeded in a 10-cm plate using growth media containing 100 U/mL penicillin and 100 μg/mL streptomycin (Gibco, ThermoFisher Scientific), 4 μg/mL blasticidin (Gibco, ThermoFisher Scientific), and 100 μg/mL hygromycin (Invitrogen, ThermoFisher Scientific). For cell lines stably expressing BioID2-FLAG, BioID2-FLAG-eGFP, BioID2-FLAG-eGFP-NLS, and BioID2-FLAG-SLX4, hygromycin resistant cells were expanded in the presence of 100 μg/mL hygromycin and pooled to create heterogenous, polyclonal stable cell lines. For cell lines expressing SLX4-FLAG-BioID2, monoclonal cells were isolated, expanded, and screened for construct expression by western blotting. Monoclonal cells that expressed the highest levels of the target construct were expanded and used for downstream analysis.

### Tetracycline-mediated expression and live cell biotinylation

Flp-In T-REx HEK293 cells were grown in complete media. Expression of the integrated construct was induced with either 1 ng/mL (BioID2-FLAG, BioID2-FLAG-eGFP, and BioID2-FLAG-eGFP-NLS) or 1 μg/mL (untransfected, BioID2-FLAG-SLX1, BioID2-FLAG-SLX4, and SLX4-FLAG-BioID2) tetracycline for 17 hr followed by the addition of biotin (50 μM final) for another 8 hr. In all cases, expression of the integrated construct was induced with tetracycline for 25 hr. The different concentrations of tetracycline were used to achieve similar protein expression across all samples.

### Immunofluorescence

Approximately 100,000 Flp-In T-REx HEK293 cells were seeded on 12 mm poly-L-lysine coated coverslips (Corning) in 12-well plates using complete growth media supplemented with biotin-depleted FBS. Cultures were grown for approximately 24 hr, after which time the cells were either left untreated, treated with tetracycline to induce expression of the integrated construct, or treated with tetracycline and biotin for live cell labeling, as described above. Cells were washed once with PBS and then permeabilized in PBS containing 0.4% Triton X-100 for 10 min at room temperature. The cells were washed once with PBS, fixed with 2% paraformaldehyde in PBS for 20 min at room temperature, and washed again in PBS. Coverslips were transferred to ice-cold methanol for 1 min and then washed twice in PBS at room temperature. The samples were blocked at room temperature in blocking buffer (PBS containing 5% goat serum and 0.05% Tween-20) for 1 hr at room temperature, transferred to a humidified chamber, and incubated overnight at 4°C with primary antibodies diluted in blocking buffer: mouse monoclonal anti-FLAG clone M2 (MilliporeSigma, AB_262044). The next day, coverslips were washed 3 x 10 min in PBS containing 0.1% Triton X-100 at room temperature and then incubated for 1 hr at room temperature with secondary antibodies diluted in blocking buffer: AlexaFluor594 goat anti-mouse IgG (Molecular Probes, ThermoFisher Scientific, AB_2534091) and AlexaFluor488 Streptavidin (Molecular Probes, ThermoFisher Scientific). All hybridized coverslips were washed 3 x 10 min in PBS containing 0.1% Triton X-100 at room temperature. Coverslips were mounted onto glass slides using Molecular Probes ProLong Gold Antifade Mountant containing DAPI (ThermoFisher Scientific) and allowed to dry at room temperature overnight before sealing with nail polish. Images were acquired on a Leica SP8 Lightning confocal laser scanning microscope equipped with a 63× oil objective and captured at 2,056 × 2,056 pixels (SickKids Imaging Facility, The Hospital for Sick Children (SickKids), Toronto). A minimum of 10 slices were taken at a Z-stack distance of 0.33 μm. Adobe Photoshop CC 2021 was used to perform minor manual adjustments for visualization.

### Preparation of cell extracts

For western blotting, pellets were prepared from Flp-In T-REx HEK293 cells grown in 15-cm plates using complete growth media containing biotin-depleted FBS. Tetracycline-mediated expression of the integrated constructs and live cell labeling with biotin were performed as described above. Cells were harvested by scraping into ice-cold PBS (137 mM NaCl, 3 mM KCl, 8.03 mM Na_2_HPO_4_, 1.83 mM KH_2_PO_4_) and pelleted by centrifugation at 500 × *g* (5 min at 4°C). Lysis was performed as described previously [39], with the following modifications. Cells were resuspended in a 2X pellet volume of RIPA lysis buffer (50 mM Tris-HCl pH 7.4, 150 mM NaCl, 10% glycerol, 1% NP-40 substitute, 0.1% SDS, 0.5% sodium deoxycholate, 1 mM EDTA) supplemented with 1 mM DTT, protease inhibitors (10 μg/mL aprotinin, 13 μM bestatin, 10 μg/mL leupeptin, 10 μg/mL pepstatin, 0.1 mM PMSF), and phosphatase inhibitors (20 mM β-glycerophosphate, 5 mM sodium fluoride, 1 mM sodium pyrophosphate, 1 mM sodium orthovanadate). The cell extracts were incubated at 4°C with gentle agitation for 20 min and then sonicated on ice using a Branson Digital Sonifier 250 with microtip probe at 10% amplitude (3 cycles of 10 s ON and 2 s OFF). Extracts were clarified by centrifugation at 16,100 × *g* for 20 min at 4°C. Protein concentration in the soluble extract was determined using the *DC* Protein Assay (Bio-Rad), as per the manufacturer’s protocol.

For small-scale affinity purification (below), pellets were resuspended in ice-cold IP lysis buffer (20 mM Tris pH 8, 100 mM NaCl, 5% glycerol, 5 mM CaCl_2_, 0.05% NP-40, 1 mM EDTA) supplemented with fresh protease inhibitors (1 mM PMSF, 10 μg/mL pepstatin A, 13 μM bestatin, 10 μg/mL leupeptin, 10 μg/mL aprotinin, 1 mM benzamide). Micrococcal nuclease was added to a final concentration of 75 U/μL. The lysates were sonicated at 4°C using a Branson Digital Sonifier 250 with microtip probe at 10% amplitude (3-5 cycles of 10 s ON and 30 s OFF). Samples were clarified by centrifugation at 16,100 × *g* for 20 min at 4°C. Protein concentration in the soluble extract was determined using the Bio-Rad Protein Assay, as per the manufacturer’s protocol. Extracts were snap-frozen in liquid N2 and stored at -80°C until affinity purification (below).

### Small-scale FLAG affinity purification

To capture FLAG-tagged proteins, 2.5 mg soluble cell extract was mixed with 37.5 μL anti-FLAG M2 agarose beads (MilliporeSigma), pre-equilibrated in IP lysis buffer, and incubated at 4°C for 3 hr with end-over-end rotation. Subsequently, samples were centrifuged at 500 × *g* (2 min at 4°C) and washed sequentially as follows: once with 1 mL IP Wash Buffer 1 (50 mM Tris-Cl pH 7.5, 150 mM, NaCl, 10% glycerol, 0.5% NP-40, 1 mM EDTA), twice with 1 mL IP Wash Buffer 2 (50 mM Tris-Cl pH 7.5, 300 mM, NaCl, 10% glycerol, 0.5% NP-40, 1 mM EDTA), and once with 1 mL IP Wash Buffer 1. Each wash involved resuspending the resin by gentle inversion, centrifuging at 500 × *g* (2 min at 4°C), and then manually removing the solution. Following the final wash, bound proteins were eluted in an equal volume of 2X sample buffer (62.5 mM Tris HCl pH 6.8, 25% glycerol, 2% SDS, 0.1% (w/v) Orange G, 50 mM DTT) and heated at 95°C for 5 min. Samples were stored at -20°C until analysis by western blotting (below).

### Western blotting

Protein samples were mixed with 4X sample buffer (125 mM Tris HCl pH 6.8, 50% glycerol, 4% SDS, 0.2% (w/v) Orange G, 100 mM DTT) to a final concentration of 1X and heated at 95°C for 10 min. Samples were resolved by SDS-PAGE using NuPAGE 4-12% Bis-Tris gels (Invitrogen, ThermoFisher Scientific) and MOPS running buffer (50 mM MOPS, 50 mM Tris base, 0.1% SDS, 1 mM EDTA), and then transferred onto 0.22 μm nitrocellulose membrane (Amersham, Cytiva) in cold transfer buffer (25 mM Tris base, 190 mM glycine, 20 % v/v methanol). Membranes were blocked for 1 hr at room temperature using Odyssey Blocking Buffer (LI-COR) and then incubated overnight (4°C) with primary antibodies diluted in Odyssey Blocking Buffer containing 0.2% Tween-20: mouse monoclonal anti-FLAG clone M2 (MilliporeSigma, AB_262044) and rabbit polyclonal anti-β-actin clone D6A8 (Cell Signalling Technology, AB_10950489). Hybridized membranes were washed three times with PBS containing 0.1% Tween-20 (10 min per wash) and then incubated for 1 hr at room temperature with LI-COR secondary antibodies diluted in Odyssey Blocking Buffer containing 0.2% Tween-20: IRDye 800CW goat anti-mouse IgG (LI-COR, AB_621842) and IRDye 680RD goat anti-rabbit IgG (LI-COR, AB_10956166). Membranes were washed in PBS containing 0.1% Tween-20 (10 min per wash) and then imaged using a LI-COR Odyssey Clx imaging platform. Biotinylated proteins were detected similarly with the following modifications. Following transfer, nitrocellulose membranes were blocked for at least 1 hr in Odyssey Blocking Buffer containing 0.2% Tween-20 and 0.1% SDS. Blocked membranes were incubated for at least 1 hr in the same solution containing IRDye 800CW Streptavidin (LI-COR), washed three times in PBS containing 0.1% Tween-20 (10 min per wash) and then imaged using a LI-COR Odyssey Clx imaging platform. All files were exported using Image Studio Lite v5.2.5 and contrasted using Adobe Photoshop CC 2021.

### Streptavidin purification of biotinylated proteins and on-bead trypsin digest

The BioID2 method was adapted from previously published work [38]. Briefly, Flp-In T-REx HEK293 cells were grown to confluency in 15-cm plates using complete growth media. Tetracycline-mediated expression of the integrated constructs and live cell labeling with biotin were performed as described above. Cells were harvested by scaping into ice-cold PBS (137 mM NaCl, 3 mM KCl, 8.03 mM Na_2_HPO_4_, 1.83 mM KH_2_PO_4_) and pelleted by centrifugation at 500 × *g* (5 min at 4°C). Cell pellets were weighed (≥ approximately 0.1 g per sample), snap frozen, and stored at -80°C.

Cells were lysed in modified RIPA buffer (50 mM Tris-HCl pH 7.5, 150 mM NaCl, 1.5 mM MgCl_2_, 1 mM EGTA, 0.1% SDS, 1% IGEPAL CA-630) supplemented with fresh sodium deoxycholate (0.4%) and Protease Inhibitor Cocktail (MilliporeSigma, P8340) at a 1:4 pellet weight:volume ratio (i.e., 400 µL buffer per 100 mg pellet). The lysate was incubated at 4°C with gentle rotation for 20 min and then sonicated at 4°C using a Qsonica Sonicator with CL-18 probe at 25% amplitude (3 cycles of 5 s ON and 3 s OFF). After sonication, the extract was treated with Benzonase (375 U per sample) and incubated at 4°C for 15 min. The sample was supplemented with SDS to bring the total concentration to 0.4% SDS, mixed gently, and incubated at 4°C for 15 min. The extract was clarified by centrifugation at 16,000 × *g* (20 min at 4°C) and the soluble fraction was transferred to a new tube.

To capture biotinylated proteins, the soluble fraction was mixed with 30 μL (bed volume) Streptavidin Sepharose HP (Cytiva) resin, pre-equilibrated in modified RIPA buffer containing 0.4% SDS, and incubated with gentle rotation at 4°C for 3 hr. Subsequently, the resin was pelleted by centrifugation at 500 × *g* (2 min at 4°C) and washed sequentially as follows: once with wash buffer (50 mM Tris-HCl pH 7.5, 2% SDS), twice with modified RIPA buffer containing 0.4% SDS, and three times with ABC buffer (50 mM ammonium bicarbonate pH 8.5). The resin was collected by centrifugation at 500 × *g* (2 min at 4°C) in between washes. After removing the ABC wash buffer, on-bead trypsin digest of peptides was performed by mixing the resin with 1 μg trypsin (MilliporeSigma) dissolved in ABC buffer and incubating at 37°C overnight with gentle rotation. The next day, an additional 0.5 μg trypsin (dissolved in ABC buffer) was added and the sample was incubated at 37°C for 2 hr, after which time the sample was gently vortexed and then centrifuged at 500 × *g* for 2 min. The supernatant was transferred to a fresh tube. The resin was washed twice with 30 μL HPLC-grade H_2_O, with intervening centrifugation at 500 × *g* for 2 min. Each wash was collected and pooled with the supernatant. The supernatant was acidified by adding 50% formic acid to a final concentration of 2% v/v, dried by vacuum centrifugation, and stored at -80°C.

### Affinity purification and on-bead trypsin digests

The FLAG affinity purification (AP)-MS protocol was adapted from [38] with minor modifications. Briefly, Flp-In T-REx HEK293 cells were grown to confluency in 15-cm plates using complete growth media. Tetracycline-mediated expression of the integrated constructs and live cell labeling with biotin were performed as described above. Cells were harvested by scraping into ice-cold PBS (137 mM NaCl, 3 mM KCl, 8.03 mM Na_2_HPO_4_, 1.83 mM KH_2_PO_4_) and pelleted by centrifugation at 500 × *g* (5 min at 4°C). Cell pellets were weighed (≥ approximately 0.1 g per sample), snap frozen, and stored at -80°C.

Cell pellets were resuspended in ice-cold AP lysis buffer (50 mM HEPES pH 8.0, 100 mM KCl, 2 mM EDTA, 0.1% NP-40 substitute, 10% glycerol) supplemented with fresh PMSF (1 mM), DTT (1 mM), and Protease Inhibitor Cocktail (MilliporeSigma, P8340) at a 1:4 pellet weight:volume ratio. The sample was subjected to 1 freeze-thaw cycles, with each cycle consisting of a 5-10 min incubation on dry ice, followed by incubation in a 37°C water bath with agitation until a small amount of ice remains. The lysate was then sonicated at 4°C using a Qsonica Sonicator with a 1/8” (3.2 mm) microtip at 30% amplitude (3 cycles of 5 s ON and 3 s OFF). After sonication, the extract was treated with TurboNuclease (250 U per sample) and RNaseA (10 μg per sample), and incubated at 4°C with end-over-end rotation for 15-20 min. The sample was clarified by centrifugation at 20,817 × *g* (20 min at 4°C) and the soluble fraction was transferred to a new tube.

To capture FLAG-tagged proteins, the soluble fraction was mixed with 12.5 μL (bed volume) anti-FLAG M2 magnetic beads (MilliporeSigma), pre-equilibrated in AP lysis buffer, and incubated at 4°C for 3 hr with end-over-end rotation. Subsequently, the sample was centrifuged at 500 × *g* (2 min at 4°C) and then placed on a magnetic rack. After removing the supernatant, the resin was washed with 1 mL AP lysis buffer, transferred to a new tube, and washed once with 1 mL AP wash buffer (20 mM Tris-HCl pH 8.0, 2 mM CaCl_2_). Each wash involved resuspending the resin by pipetting up and down 4 times in buffer, placing the tube on the magnetic rack for 30 s, and then removing the buffer. After the final wash, on-bead trypsin digest of peptides was performed by resuspending the resin in 7.5 μL trypsin digestion buffer (100 ng/μL trypsin in 20 mM Tris-HCl pH 8.0) and incubating overnight at 37°C with gentle agitation. The next day, the sample was centrifuged at 500 × *g* for 1 min at room temperature and then magnetized for 30 s. The supernatant was transferred to a new tube. The resin was resuspended in an additional 2.5 μL trypsin digestion buffer and incubated for 3 hr at 37°C (no agitation), after which time the supernatant was collected as described above. The supernatant was acidified by adding 50% formic acid to a final concentration of approximately 5% v/v and stored at -20°C until analysis by mass spectrometry.

### Mass spectrometry analysis

Affinity purified and digested peptides were analyzed using data-dependent acquisition (DDA) nanoscale high performance liquid chromatography (nano-HPLC) coupled to tandem mass spectrometry (MS/MS). One-sixth of the BioID2 samples and one-quarter of the AP samples were used for analysis. Nano-spray emitters were generated from fused silica capillary tubing (100 µm internal diameter, 365 µm outer diameter, 5-8 µm tip opening) using a laser puller (Sutter Instrument Co., model P-2000), with parameters set as follows: heat = 280, FIL = 0, VEL = 18, and DEL = 2000. Nano-spray emitters were packed with C18 reversed-phase material (Reprosil-Pur 120 C18-AQ, 3 µm) resuspended in methanol using a pressure injection cell. Samples in 5% formic acid were directly loaded onto a 100 µm × 15 cm nano-spray emitter at 800 nL/min for 20 min. Peptides were eluted from the column using a linear acetonitrile gradient generated by an Ekspert nanoLC 425 (Eksigent, Dublin CA) and analyzed on a TripleTOF 6600 instrument (AB SCIEX, Concord, Ontario, Canada). The 90 min gradient was delivered at 400 nL/min from 2% acetonitrile with 0.1% formic acid to 35% acetonitrile with 0.1% formic acid. This was followed by a 15 min wash using 80% acetonitrile with 0.1% formic acid and a 15 min equilibration in 2% acetonitrile with 0.1% formic acid. The first DDA scan had an accumulation time of 250 ms and a mass range of 400 – 1,800 Da. This was followed by 10 MS/MS scans of the top 10 peptides identified in the first DDA scan, with an accumulation time of 100 ms and a mass range of 100 – 1800 Da for each MS/MS scan. Candidate ions were required to have a charge state of 2+ to 5+ and a minimum threshold of 300 cps, isolated using a window of 50 mDa. Previously analyzed candidate ions were dynamically excluded for 7 s.

### Mass spectrometry data search

All mass spectrometry data files were stored, searched, and analyzed using ProHits Laboratory Information Management System (LIMS) platform [52]. Within ProHits, WIFF files were converted to an MGF format using the WIFF2MGF converter and to an mzML format using ProteoWizard (V3.0.10702) and the AB SCIEX MS Data Converter (V1.3 beta). The data was searched using Mascot V2.3.02 [52] and Comet V2016.01 rev.2 [53]. The spectra were searched with the human and adenovirus sequences in the NCBI Reference Sequence Database (version 57, January 30^th^, 2013), supplemented with “common contaminants” from the Max Planck Institute (http://maxquant.org/contaminants.zip) and the Global Proteome Machine (GPM; ftp://ftp.thegpm.org/fasta/cRAP/crap.fasta), forward and reverse sequences (labeled “gi|9999” or “DECOY”), sequence tags, and streptavidin, for a total of 72,482 entries. Database parameters were set to search for tryptic cleavages, allowing up to 2 missed cleavages per peptide with a mass tolerance of 35 ppm for precursors with charges of 2+ to 4+ and a tolerance of 0.15 amu for fragment ions. Variable modifications were selected for deamidated asparagine and glutamine, and oxidated methionine. Results from each search engine were analyzed through TPP (the Trans-Proteomic Pipeline, v.4.7 POLAR VORTEX rev 1) via the iProphet pipeline [54]. All proteins with an iProphet probability ≥ 95% and 2 unique peptides were used for analysis.

### Protein-protein interaction scoring

Significance analysis of interactome express (SAINTexpress) version 3.6.1 was used to calculate the probability of potential protein-protein associations/interactions compared to background contaminants using default parameters [55]. In brief, SAINTexpress is a statistical tool that compares the spectral counts of each prey identified with a given BioID2 bait against a set of negative controls. For BioID, negative controls consisted of streptavidin affinity purifications from untransfected cells and cells expressing BioID2-FLAG, BioID2-FLAG-eGFP, and BioID2-FLAG-eGFP-NLS (three biological replicates each). For AP-MS, negative controls consisted of anti-FLAG affinity purifications from BioID2-FLAG, BioID2-FLAG-eGFP, and BioID2-FLAG-eGFP-NLS (three biological replicates each). Three biological replicates were collected for all cell lines and conditions. For BioID protein-protein interaction scoring, two replicates with the highest spectral counts for each prey were used for baits; three replicates were used for negative controls for SAINTexpress. For AP-MS protein-protein interaction scoring, three replicates were used for baits and negative controls for SAINTexpress. SAINT scores were averaged across replicates and these averages were used to calculate a Bayesian False Discovery Rate (BFDR); preys with BFDR ≤ 1% were considered high-confidence protein interactions. All non-human protein interactors (did not start with “NP” in Prey column) were removed from the SAINT analysis, except for BirA_R118G_H0QFJ5.

### Cytoscape analysis

Proteins detected in at least two biological replicates and with a BFDR ≤ 1% were considered high-confidence interactions. Cytoscape v3.9.1 [56] was used to create network figures for BioID and AP-MS results for SLX1 and SLX4. The size of each protein “node” is proportional to the unique peptide count for each protein, averaged across all biological replicates. Node size for each dataset was dependent on detected proteins, where the protein with the highest average unique peptide count in each dataset was assigned the largest node size. Node color is related to the spectral count of each protein, averaged across all biological replicates. Spectral count is a measure of how many times a protein was detected by mass spectrometry and serves as a measure of confidence for each hit.

### Gene ontology enrichment analysis

Gene ontology (GO) enrichment analysis was performed using the high-confidence interactors as inputs. The Princeton Generic Gene Ontology (GO) Term Finder was used for the enrichment analysis [57]. Default options were used to view biological processes in *Homo sapiens.* GO enrichment results were inputted into Revigo [58] to remove redundant GO terms. Resulting list size is indicated in the figure legends. All terms that correspond to ≥ 10% genome frequency were manually removed.

### Data Visualization

The STRING database was used to model the SLX4 interaction network using the high-confidence candidate partners detected by either AP-MS or BioID in at least two biological replicates. Cytoscape V3.9.1 [56] was used to visualize the network with a confidence score cut-off of 0.70. STRING interactions are scored based on text mining of literature, computational predictions from co-expression and conserved genomic context, as well as databases with interaction experiments and annotated complexes and pathways. The width of edges, or lines connecting protein nodes, correspond to STRING scores, with thicker lines denoting a higher score and thus greater confidence in the interaction. The ClusterONE application (Nepusz et al., 2012) was used to group proteins into functional complexes. Shapes and colors correspond to the STRING classification of each protein, where filled circles represent high significance nodes and filled rectangles represent nodes with multiple clusters (overlap). Grey circles and diamonds represent the least significant nodes and outliers (unclustered interactions), respectively. White circles represent nodes that did not pass the confidence cut-off score.

### Data and software availability

The BioID proteomics data have been deposited as a complete submission to the MassIVE repository and assigned the accession number MSV000090338. The ProteomeXchange accession is PXD036769. The BioID dataset is currently available for reviewers at ftp://MSV000090338@massive.ucsd.edu. Please login with username MSV000090338_reviewer; password: SLX4. The datasets will be made public upon acceptance of the manuscript.

The AP-MS proteomics data have been deposited as a complete submission to the MassIVE repository and assigned the accession number MSV000090314. The ProteomeXchange accession is PXD036690. The AP-MS dataset is currently available for reviewers at ftp://MSV000090314@massive.ucsd.edu. Please login with username MSV000090314_reviewer; password: SLX4. The datasets will be made public upon acceptance of the manuscript.

## RESULTS

### Expression and Validation of BioID2 Fusion Proteins

Human SLX4 was fused to a mutated biotin ligase from *A. aeolicus* (denoted here as BioID2) [43] and a FLAG tag at either the N- or C-terminus, resulting in two expression constructs (e.g., BioID2-FLAG-SLX4 and SLX4-FLAG-BioID2, respectively). The SLX4 constructs were stably integrated into Flp-In T-REx HEK293 cells, allowing for tetracycline-inducible expression. As controls, we generated cell lines that conditionally express BioID2-FLAG, BioID2-FLAG-eGFP, and BioID2-FLAG-eGFP fused to a nuclear localization signal (NLS) (i.e., BioID2-FLAG-eGFP-NLS). Untransfected Flp-In T-REx HEK293 cells were included as an additional control for non-specific binding and biotinylation.

We validated protein expression by immunoblotting lysates from cells treated with and without tetracycline with anti-FLAG antibodies (Figure S1B). Preliminary experiments showed that the control constructs were expressed to a significantly higher level than the SLX4 constructs (data not shown). We attempted to normalize protein expression by reducing the concentration of tetracycline used to induce the expression of the control constructs. This approach was successful, although the BioID2-FLAG-eGFP-NLS and BioID2-FLAG-eGFP proteins were still more highly expressed than the SLX4 constructs (Figure S1B). We also observed that the N-terminally tagged SLX4 construct was reproducibly expressed to higher levels compared to the C-terminally tagged counterpart (Figure S1B).

To test for live cell biotinylation, HEK293 Flp-In T-REx cells were treated with tetracycline for 17 hr and then supplemented with 50 μM biotin for 8 hr (25 hr induction with tetracycline). Cell lysates were probed with streptavidin-IRDye800 to visualize biotinylated proteins (Figure S1B). Biotinylation by BioID2-FLAG-eGFP-NLS and BioID2-FLAG-eGFP was evident by multiple proteins that bound streptavidin-IRDye800. Similar results were observed for BioID2-FLAG-SLX4 and SLX4-FLAG-BioID2, where the relative levels of biotinylation mirrored the expression of the baits (Figure S1B). The low levels of biotinylation observed in lysates from cells treated with tetracycline alone reflects the presence of biotin in cell growth medium, demonstrating the sensitivity of live cell labeling (Figure S1B).

The subcellular localization of BioID2-FLAG-tagged SLX4 proteins were investigated *in situ* using immunofluorescence confocal microscopy with anti-FLAG antibodies (Figures 1B-E). In parallel, we stained for biotinylation using streptavidin-Alexa Fluor 488. Untransfected cells were included to gauge non-specific antibody staining, and cells expressing BioID2-eGFP-NLS were analyzed to assess nuclear localization (Figures 1B, 1C). As expected, BioID2-eGFP-NLS was enriched in the nucleus (Figure 1C). When cells expressing BioID2-eGFP-NLS were supplemented with biotin, we observed intense staining for biotinylated proteins in the nucleus, some of which resided in DAPI-poor regions that likely represent nucleoli (Figure 1C).

Previous studies showed that the SLX1-SLX4 heterodimer is enriched in the nucleus, consistent with its role in DNA repair and recombination [59, 60]. Moreover, over-expressed SLX4 forms subnuclear foci in cells under basal growth conditions, and some of these foci correspond to telomeres [14, 24, 25, 61]. Likewise, BioID2-FLAG-SLX4 and SLX4-FLAG-BioID2 formed numerous discrete foci throughout the nucleus upon tetracycline addition (Figures 1D, 1E). We observed considerable spatial overlap between FLAG- and streptavidin-positive foci when cells were treated with tetracycline and biotin, indicative of live cell biotinylation of the bait and proteins in close proximity of the bait (Figures 1D, 1E).

Lastly, we performed a small-scale co-immunoprecipitation experiment to validate that the BioID2-FLAG tag did not disrupt the ability of SLX4 to interact with the SLX1 or XPF endonucleases. We chose these protein partners because they interact with distinct regions of the SLX4 scaffold (Figure 1A) [17]. Specifically, SLX1 binds the conserved C-terminal domain (CCD) of SLX4, spanning residues 1633-1834. XPF interacts with a region in the SLX4 N-terminus called the MUS312-MEI9 interaction-like region (MLR), which spans amino acid resides 409-555. Western blots showed that BioID2-FLAG-SLX4 and SLX4-BioID2-FLAG pulled down SLX1 and XPF, indicating that the BioID2-FLAG tag does not interfere with these protein-protein interactions (Figure 1F). Together, these results indicate that the BioID2-FLAG tag does not cause significant structural changes to SLX4.

### Identification of Candidate SLX4 Interacting Proteins by BioID and AP-MS

Having confirmed that the BioID2-FLAG tag did not interfere with the subcellular localization or select binding partners of SLX4, we conducted three large-scale BioID2 and FLAG affinity purifications and used mass spectrometry to identify the proteins associated with each of the bait constructs. In the BioID2 approach, biotinylated proteins are isolated using streptavidin agarose beads (Figure S1A). In contrast, during FLAG affinity purification (AP), FLAG-tagged proteins (e.g., BioID2-FLAG-SLX4) and their interacting partners are recognized and bound by anti-FLAG M2-conjugated agarose beads (Figure S1A). A difference that explains the relative orthogonality of the two techniques is that BioID uses harsh lysis and purification conditions to attempt to capture only biotinylated proteins [39]. In contrast, AP-MS uses gentle lysis and purification conditions to preserve protein complexes.

We treated cells with tetracycline for 25 hr in each approach to induce expression of the integrated BioID2-FLAG-tagged bait construct. For the BioID2 experiments, we added exogenous biotin (50 μM) to the cell cultures at 17 hr post-induction and performed live cell labeling for 8 hr. We included several controls in these experiments to obtain high-quality and high-confidence datasets: untransfected Flp-In T-REx HEK293 cells (BioID), BioID2-FLAG (BioID and AP-MS), BioID2-FLAG-eGFP-NLS (BioID and AP-MS), and BioID2-FLAG-eGFP (BioID and AP-MS).

We used SAINT*express* to assess the confidence of each bait-prey interaction detected in at least two biological replicates [55]. SAINT*express* is a newer implementation of the Significance Analysis of INTeractome (SAINT) computational tool, which uses spectral counts in the experimental and control conditions to predict the probability of a true interaction for each bait-prey pair [62]. Prey proteins were filtered based on the Bayesian False Discovery Rate (BFDR), where only proteins captured in at least two biological replicates with a BFDR ≤ 1% were considered for further analysis.

This yielded 156 and 81 high-confidence hits for the SLX4 BioID and AP-MS datasets, respectively (Figure 2A). To gauge the performance of these assays, we determined the number of preys that represent known interactors, which we defined as SLX4 binding proteins detected in at least one low-throughput and/or at least one high-throughput experiment, and as reported in BioGRID (Table 1) [63]. Based on these criteria, the BioID and AP-MS datasets each contained 50% (or 12/25) of the known SLX4 binding proteins, representing 7% and 15% of the high-confidence prey proteins, respectively (Figures 2B, C). Importantly, this list included proteins representing constitutive SLX4 binders, namely SLX1, XPF-ERCC1, and SLX4IP, as well as proteins recruited transiently to the SLX4 scaffold, including PLK1 and MUS81-EME1 (Figure 2A, Table 1). We also detected proteins with functions in transcription and transcriptional regulation (i.e., PAF, ELL, CDC73, and GTF2F), chromatin remodeling (i.e., CREBBP), and post-translational modifications (i.e., PGAM5 and CDC7). This analysis revealed one uncharacterized protein, DHX40, which is predicted to be a member of the DExH/D box family of ATP-dependent RNA helicases that have essential roles in RNA metabolism. The percentage of overlapping proteins is similar to other comparisons of AP-MS and BioID datasets [38, 47]. Most SLX4 interactors captured by both methods are novel protein partners (Figure 2B, 2C).

**Figure 2.**
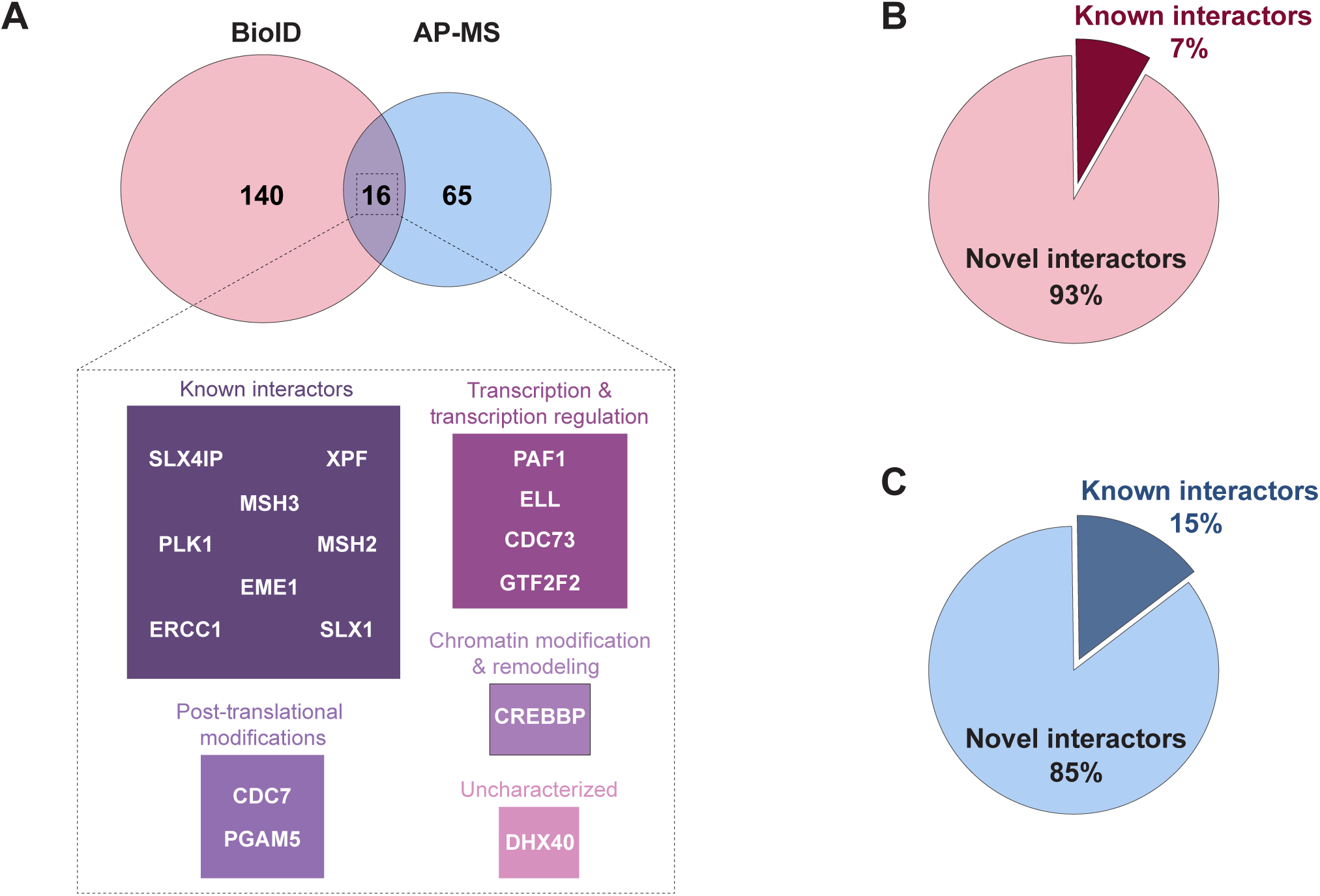
BioID and AP-MS reveal known and novel SLX4 binding partners. **(A)** Comparison of the total number of high-confidence preys that appeared in at least two biological replicates of BioID (left), AP-MS (right), or both (middle) with a Bayesian False Discovery Rate (BFDR) of ≤ 0.01. A detailed summary of the overlapping candidates categorized by protein function is shown below the Venn diagrams. **(B-C)** Approximation of previously identified SLX4 interactors *versus* novel protein partners detected by BioID **(B)** or AP-MS **(C)**.

**Table 1.** List of known SLX4 binding proteins. We defined the known SLX4 binding proteins as those that have been identified in at least one low-throughput method and/or at least one high-throughput method (asterisks), as indicated on BioGRID [56]. Brackets indicate known binding partners that were not detected by either BioID or AP-MS in this study.

| **Known SLX4 Binding Proteins** |  |
| --- | --- |
| *Interactors that meet both criteria\** | *Interactors that meet one criterion\** |
| EME1 | BRCA1 |
| ERCC1 | [FANCD2] |
| MSH2 | MRE11A |
| MSH6 | MSH3 |
| MUS81 | NBN |
| PLK1 | [PCNA] |
| SLX1A | PGAM5 |
| SLX4IP | PIAS4 |
| [SUMO2] | SUMO1 |
| RTEL1 | [SUMO3] |
| TERF2 |  |
| TERF2IP |  |
| TOPBP1 |  |
| XPF |  |

### Network and Gene Ontology Analysis of the SLX4 Interactomes

To gain an overview of the cellular localization of the SLX4 interactome, we performed a gene ontology (GO) analysis based on the cellular component for the high-confidence hits for each prey. This analysis showed that the preys were primarily associated with the nucleus and nuclear compartments (e.g., chromatin, telomeres, nuclear bodies), consistent with the well-established roles of the SLX4 scaffold in DNA repair and chromosome stability (Figure 3A, S2).

**Figure 3.**
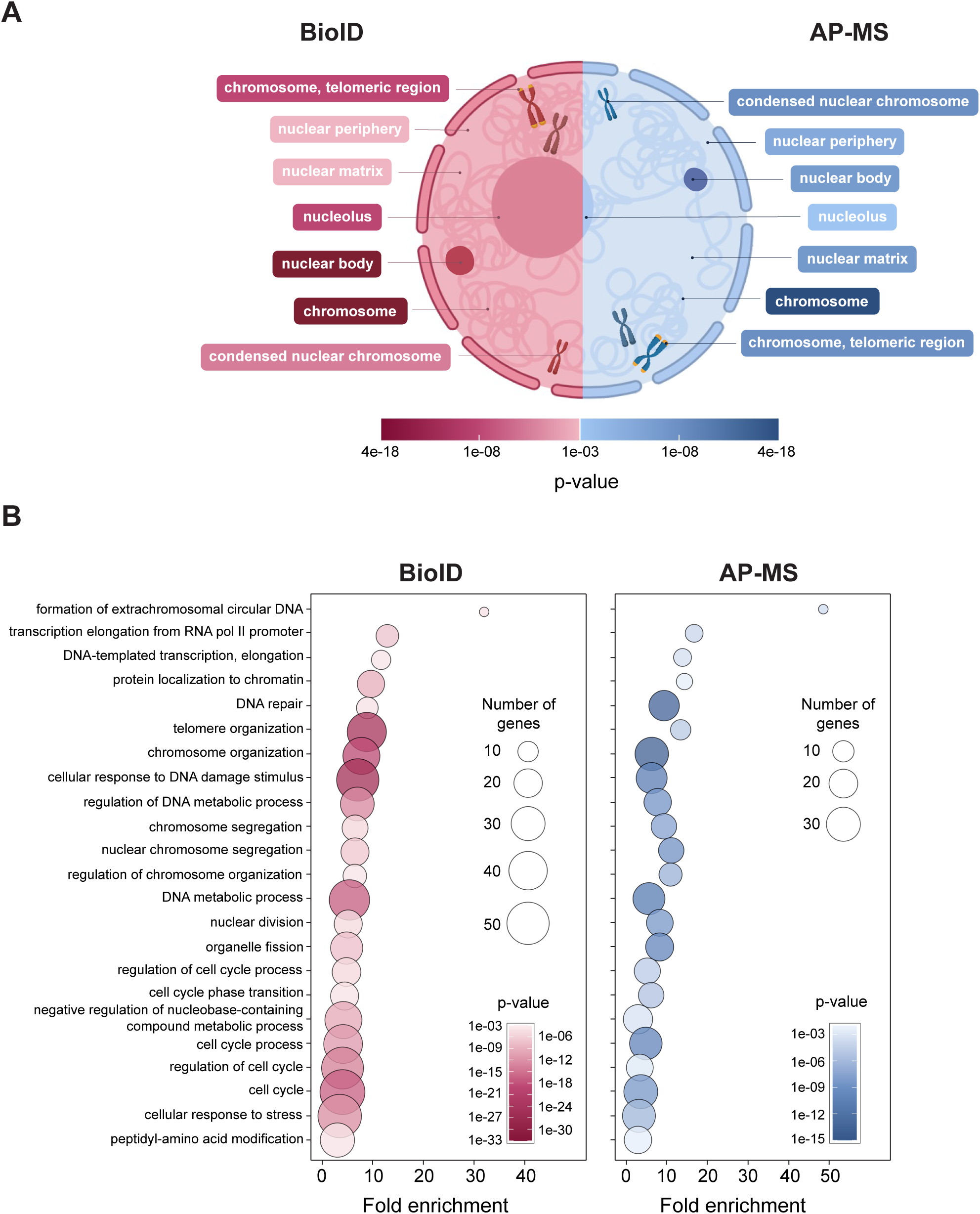
Gene ontology (GO) analysis of high-confidence SLX4 interacting proteins detected by BioID and AP-MS. **(A)** Cellular component GO terms associated with SLX4 protein hits captured by BioID (red) or AP-MS (blue). Terms were selected from Revigo after applying a small list cut-off (0.5) [58]. P-values as listed by Princeton Generic GO Term finder [57]. **(B)** Bubble plot displaying biological function GO terms associated with SLX4 binding proteins captured by BioID (red) or AP-MS (blue). Terms were selected from Revigo after applying a medium list cut-off (0.7) [58]. P-values as listed by Princeton Generic GO Term finder [57]. Fold enrichment represents the number of genes associated with a specific GO function in each proteomics dataset compared to the number of genes related to that function in the human genome.

We next aimed to identify biological pathways or functions within the SLX4 interactome. To this end, we performed a GO analysis of the biological processes associated with the SLX4 datasets (Figure 3B). GO analysis was performed using the Princeton Generic GO Term Finder [57] and refined with Revigo [58] to remove redundant terms. Proteins associated with DNA damage and cellular stress responses were enriched across both AP-MS and BioID results. We observed the same trend for proteins that function in DNA replication, chromosome segregation, and cell cycle pathways, which aligns with the known roles of the SLX4 scaffold. Nevertheless, some biological processes were more enriched in the AP-MS dataset, including the formation of extrachromosomal circular DNA, chromosome organization, and nuclear chromosome segregation. These results could reflect long-distance protein interactions that are outside the BioID2 biotinylation radius or lack surface-exposed lysine residues for biotinylation. We observed a strong enrichment of proteins involved in transcription elongation in both datasets (Figure 3B). These results likely indicate a role for SLX4 in resolving replication-transcription conflicts [64] or other DNA lesions to promote transcription elongation.

To facilitate direct comparison between the preys identified in the SLX4 BioID and AP-MS datasets, we arranged the high-confidence hits into interaction networks using Cytoscape (Figure 4A, B) [56]. For each network, node size and color reflect the confidence in each protein-protein interaction. Node color indicates average spectral count, where a darker color signifies a higher spectral count and thus higher confidence in an interaction. Node size denotes the average unique peptide count, with a larger node designating a higher unique peptide count and greater confidence in an association. Notably, proteins detected by AP-MS have higher spectral and unique peptide counts than those detected by BioID, implying more robust interactions with SLX4 (Figure 4A, B). Proteins in each network were manually grouped based on biological function.

**Figure 4.**
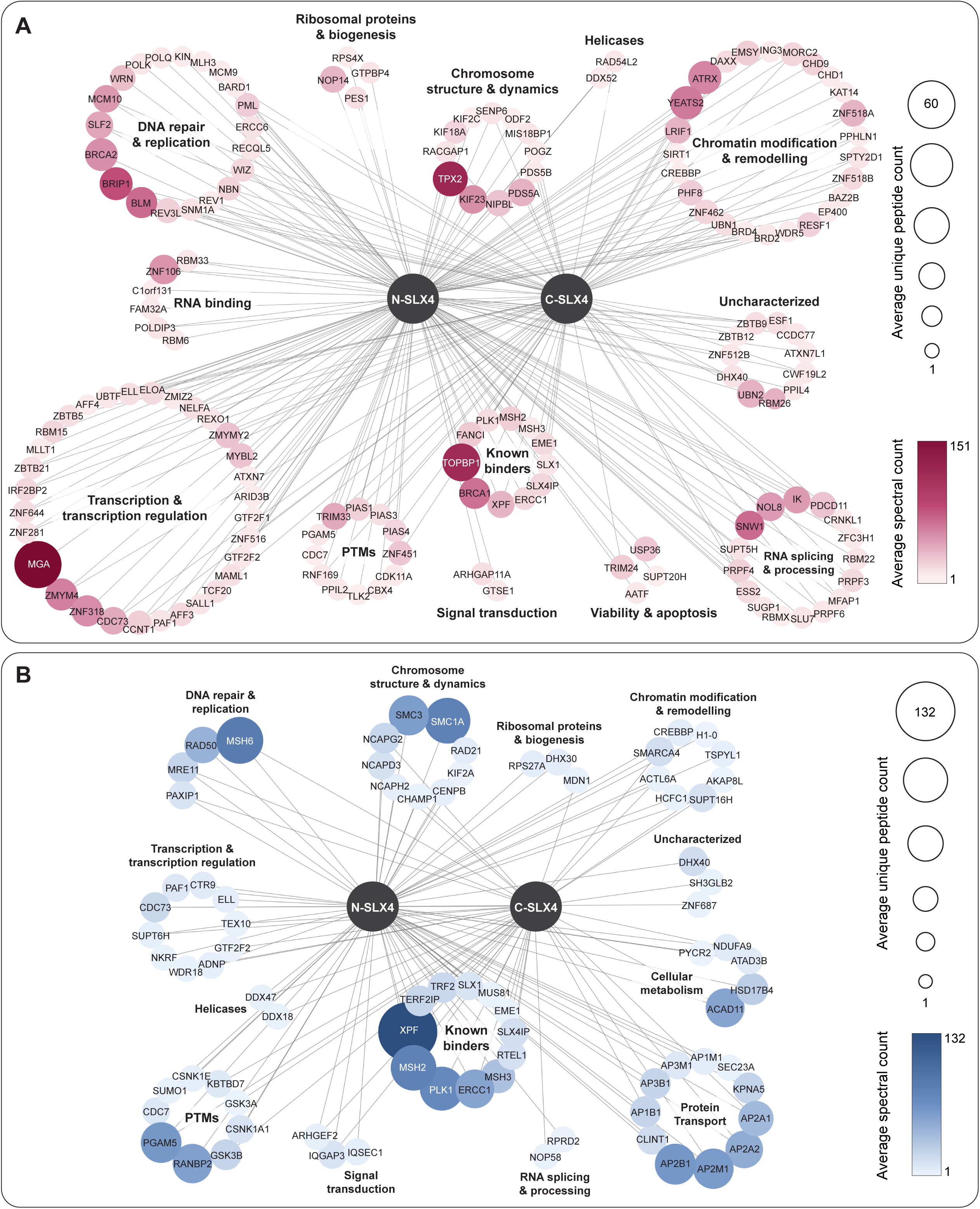
Network maps of SLX4 binding proteins identified by BioID and AP-MS. **(A-B)** High-confidence SLX4 interactors detected in at least two biological replicates of BioID **(A)** or AP-MS **(B)**. Proteins (nodes) were manually clustered in Cytoscape based on biological function [56]. Spectral count is a measure of the number of times a specific protein is detected by mass spectrometry. Unique peptide count reflects the number of peptides detected by mass spectrometry that correspond to one protein. Values were averaged over 2 or 3 biological replicates and are represented according to the scale bars found to the right of each network.

The wide array of functional categories highlights the diversity of SLX4 protein partners. High-confidence hits captured by both techniques include known SLX4 partners, including XPF-ERCC1, SLX4IP, MSH2-MSH3, and TOPBP1, lending credence to the overall results (Figure 4A, B) [10, 12–15, 27, 30, 31, 33, 37, 65]. Several high-confidence hits were associated with DNA repair and replication, consistent with the well-established roles of SLX4 in homologous recombination. We captured these protein-protein interactions without exogenous DNA-damaging agents, reflecting the basal DNA repair processes that occur during unperturbed cell growth and highlighting the robustness of our approach.

The number of functional categories was similar in both datasets (Figure 4A, B). However, the BioID dataset was noticeably more enriched in proteins with roles in DNA repair and replication (20/156 [13%] *vs.* 4/81 [5%]), RNA splicing and processing (16/156 [10%] *vs.* 2/81 [2%]), and transcription or transcription regulation (31/156 [20%] *vs.* 10/81 [12%]). Reciprocally, the AP-MS dataset uniquely captured proteins with roles in metabolism and protein transport (Figure 4B). As each method has its strengths and weaknesses, our work highlights the value of using proximity labeling and affinity purification to gain detailed insight into the binding partners of a target protein. This may be especially pertinent when the target protein fulfills a scaffold function, as for SLX4. For example, our results indicate that SLX4 is in close spatial proximity to factors that function in RNA splicing and processing, which could reveal an understudied function of SLX4 in RNA metabolism.

### Compilation of a Comprehensive SLX4 Interactome

Driven by our motivation to define a comprehensive SLX4 interactome, we combined the BioID and AP-MS datasets (237 high-confidence preys) and used the ClusterONE Cytoscape plug-in to identify densely connected regions that usually represent multiprotein complexes or subcomplexes [66]. Select proteins were grouped into clusters based on their cohesiveness or degree of interconnection (Figure 5). From the global SLX4 interactome, 12 clusters were identified (8 clusters with p < 0.05), although proteins within the clusters often demonstrated functional overlap (Figure 5). The clusters were generally associated with the following biological processes: (i) DNA repair (35 proteins, p < 0.05, green), (ii) transcription (15 proteins, p < 0.05, hot pink), (iii) ribosome and RNA biogenesis (14 proteins, p < 0.05, magenta pink), (iv) chromosome structure and dynamics (12 proteins, p < 0.05, lemon), (v) chromatin remodeling (12 proteins, p < 0.05, dark orange), (vi) RNA processing (11 proteins, p < 0.05, red), (vii) chromosome cohesion and condensation (10 proteins, yellow orange), (viii) protein transport (9 proteins, p < 0.05, cobalt blue), (viv) protein sumoylation (8 proteins, light blue), (x) kinetochore assembly (4 proteins, lilac), (xi) protein phosphorylation (4 proteins, p < 0.05, teal), and (xii) protein ubiquitylation (3 proteins, magenta).

**Figure 5.**
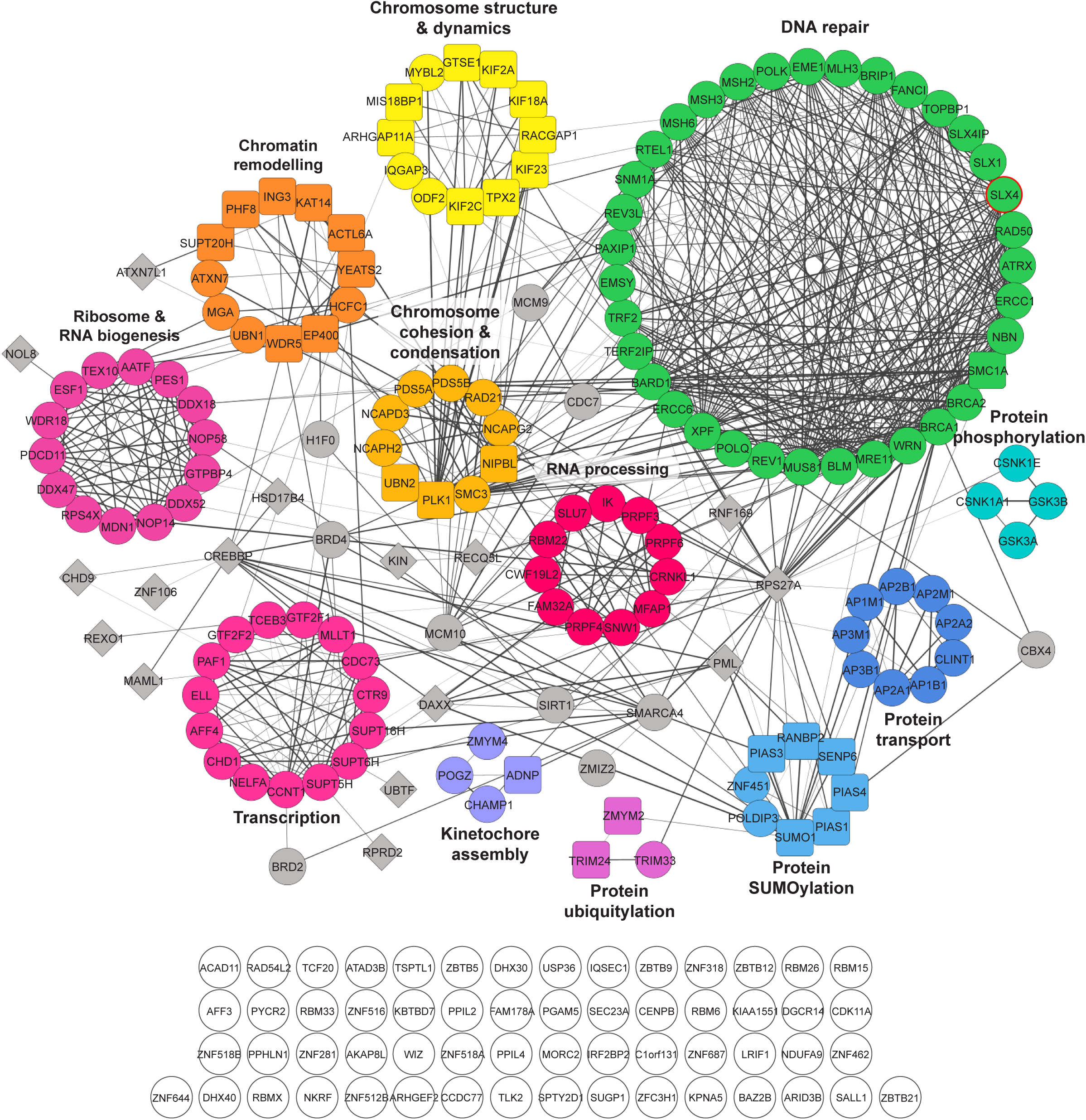
Compilation of a comprehensive SLX4 interaction network using proximity labeling and affinity purification proteomics. SLX4 protein interaction network based on preys identified in at least two biological replicates of BioID and AP-MS. Proteins (nodes) were clustered using Cytoscape ClusterONE, which uses the STRING database to group nodes into functional complexes; the confidence score cut-off was set to 0.70 (high-confidence) [107]. Line (edge) thickness corresponds to STRING scores, with thicker edges representing higher confidence interactions between nodes. Shapes and colors correspond to the STRING classification of each protein, where filled circles represent high significance nodes and filled rectangles represent nodes with multiple clusters (overlap). Grey circles and diamonds represent the least significant nodes and outliers (unclustered interactions), respectively. White circles represent nodes that did not pass the confidence cut-off score. The SLX4 node is outlined in red.

In a separate analysis, we manually grouped the high-confidence preys (237 in total) by biological function (Figure 6). This approach increased the scope of biological functions, thus permitting an in-depth examination of the diversity of the SLX4 interactome. We assigned 22 biological functions, which include both expected and unexpected cell processes (Figure 6, see Discussion). The expected or canonical functions include DNA repair and replication (36/237 preys; 15%), chromatin remodeling and modification (30/237 preys; 13%), chromosome structure and dynamics (17/237 preys; 7%), telomere function and maintenance (3/237 preys; 1%), chromosome condensation (3/237 preys; 1%), and several post-translational modification enzymes (i.e., kinases, phosphatases, sumoylation, and ubiquitylation; 17/237 preys; 7%). Altogether, 45% of the high-confidence preys were associated with biological processes that are in line with the best-characterized roles of the SLX4 scaffold in genome stability maintenance [17].

On the other hand, 55% of the high-confidence preys were associated with biological processes that have poorly characterized or emerging roles for the SLX4 scaffold (Figure 6, see Discussion). For example, we observed several potential links between SLX4 and RNA splicing and processing (18/237 preys; 8%), transcription and transcriptional regulation (37/237 preys; 7%), and RNA binding proteins (5/218 preys; 2%). We also observed unanticipated links between SLX4 and biological pathways related to cellular metabolism (6/237 preys; 3%), signal transduction (5/237 preys; 2%), and others (Figure 6). Additionally, 6% of the high-confidence preys are proteins with uncharacterized functions. That we observed fewer high-confidence hits with each of these biological processes suggests that they may be more likely to represent false-positive interactions. This may be of particular concern for the interactions that were detected uniquely by AP-MS and could reflect protein-protein associations that arise during lysis (e.g., protein trafficking and transport, clathrin-dependent endocytosis). Further work is needed to establish the functional relevance of these unexpected associations. Despite these potential caveats, our study reveals the biological diversity of the SLX4 scaffold and uncovers many exciting avenues for future research.

**Figure 6.**
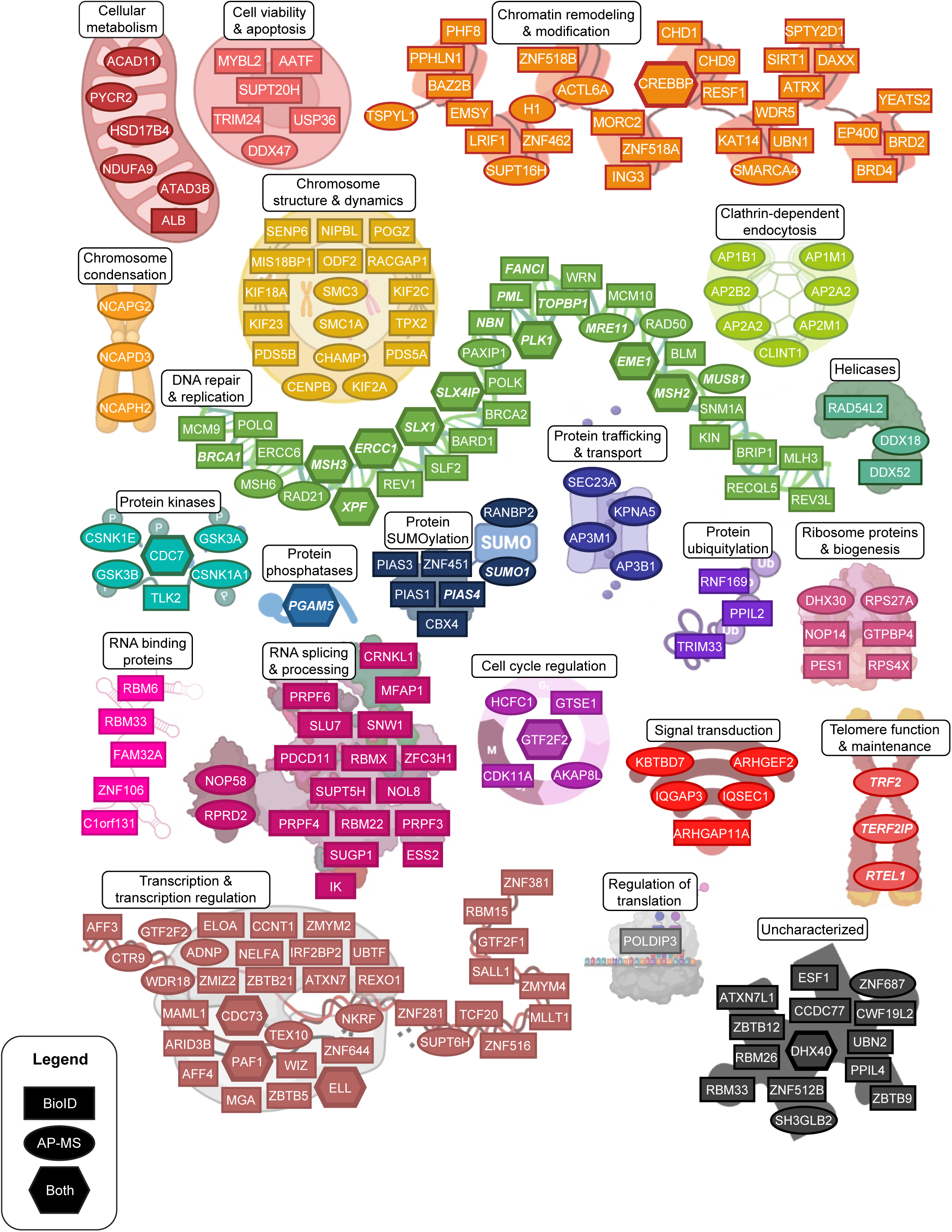
Functional landscape of the SLX4 interaction network. Candidate SLX4 binding partners identified in at least two biological replicates of BioID and AP-MS were grouped based by function. Shapes indicate the method(s) used to identify high-confidence preys, as defined in the legend (i.e., BioID2, AP-MS, or both). Known SLX4 interactors are bolded and italicized.

## DISCUSSION

The DNA repair scaffold SLX4 has multi-faceted roles in DNA repair, which reflect its interactions with various partner proteins. However, a comprehensive and unified analysis of the SLX4 interactome is lacking. We used BioID and AP-MS to fill this gap and explore the complete repertoire of SLX4 binding partners in human cells. While our studies identified many known SLX4 interactors, most of our high-confidence hits represent novel SLX4 binding proteins (Figure 2, Table 1). Given that insight into protein function often results from a better understanding of protein partners, our datasets provide a wealth of new insight into the functional landscape of the SLX4 scaffold.

The presence of known SLX4 binding proteins in the BioID and AP-MS datasets alleviates potential concerns that the BioID2-FLAG tag could compromise the nuclear localization or binding partners of SLX4 (Table 1). For example, we detected SLX1 and XPF-ERCC1 in both datasets, representing two well-known structure-selective endonucleases that bind the SLX4 scaffold (Figure 1A). We also detected MUS81-EME1, a structure-selective endonuclease that is transiently recruited to the SLX4 scaffold in early mitosis [10, 15, 20]. This result highlights the presence of temporally-regulated protein partners in our datasets, as does the interaction between SLX4 and PLK1. We further identified the association between SLX4 and the MSH2-MSH3 mismatch recognition protein, recently shown to stimulate Holliday junction resolution by the SMX complex [30]. Importantly, we captured interactions between SLX4 and several proteins that could fulfill a recruitment role and deliver SLX4 to telomeres (TRF2, TERF2) [24, 25] and stalled or collapsed replication forks (RTEL1, FANCI, TOPBP1) [64, 65, 67].

Human SLX4 is a large protein that is predicted to contain extensive regions of disorder (Figure 1A) [17]. As such, SLX4 likely adopts a range of extended conformations instead of a compact globular structure. These spatial considerations warrant special attention in the experimental design of proximity-dependent labeling experiments. As such, we generated SLX4 bait constructs with the BioID2-FLAG tag fused to the SLX4 N- or C-terminus. We reasoned that this approach would minimize the number of false negatives in our proximity-dependent labeling experiments (i.e., prey is not detected because it is beyond the 10 nm labeling distance), thus maximizing the number of SLX4 interactors detected by BioID. For example, SLX1 interacts with the conserved C-terminal domain (CCD) of SLX4 (Figure 1A) and, indeed, was only detected as a binding partner of SLX4-FLAG-BioID2 (Figure 4A). These results underscore the benefit of using N- and C-terminally tagged prey constructs in our proximity-dependent labeling experiments. Conversely, when analyzed by AP-MS, SLX1 appeared as a high-confidence interactor of BioID2-FLAG-SLX4 and SLX4-FLAG-BioID2 because this method does not depend on biotinylation. Together, these results highlight the value of combining BioID and AP-MS to generate protein-protein interaction networks.

Although we identified many known SLX4 binding proteins, a few were missing from one or both datasets (Table 1). Preys missing from both datasets could be due to different cell types, low abundance of certain interactors or interactions, the rapid reversibility of some protein-protein interactions, or inefficient tryptic digests. Since we collected samples from asynchronous cell cultures, transient or conditional interactions within the short subphases of mitosis (e.g., metaphase, anaphase, telophase) or cytokinesis could be missed. Another possibility is that cells need to be treated with genotoxic agents to enrich for interactions between SLX4 and known DNA repair proteins. Some SLX4 binding proteins were only captured by one proteomics method. For example, TOPBP1 was only detected by BioID, potentially reflecting a transient protein-protein interaction. On the other hand, MUS81 was only detected by AP-MS. This may be due to the lack of surface-exposed lysine residues, a prerequisite for biotinylation. These and other examples further highlight the value of using BioID and AP-MS to determine protein networks.

Our ultimate goal in this study was to provide a comprehensive interaction map of the human SLX4 scaffold. To this end, we used a systematic approach that first involved grouping the high-confidence preys from each technique into interaction networks based on function (Figure 4), followed by the compilation and organization of all high-confidence preys into functional clusters (Figures 5, 6). These detailed functional groupings underscore the value of combining BioID and AP-MS, as outlined above. For example, preys involved in RNA metabolism and chromatin biology were almost entirely uniquely detected by BioID (Figure 6). SLX4 interactors that catalyze post-translational modifications, such as ubiquitylation and sumoylation, were also skewed towards detection by BioID. Conversely, preys involved in chromosome condensation, protein transport, and clathrin-dependent endocytosis were exclusively identified by AP-MS (Figure 6). Importantly, DNA repair and replication proteins were identified in both proteomics methods, likely because these represent the most robust SLX4 binding partners (e.g., SLX1, XPF-ERCC1, SLX4IP). Similarly, proteins with a role in chromosome structure and dynamics were detected in BioID and AP-MS (Figure 6). Select examples of biological pathways and protein complexes are discussed in depth below.

### DNA Repair: The Fanconi Anemia Pathway

Fanconi anemia is a rare genetic disorder characterized by physical abnormalities (e.g., short stature, irregular skin coloring, skeletal malformations), bone marrow failure, and an increased risk of certain malignancies (e.g., acute myeloid leukemia, head and neck squamous cell carcinomas) [68]. At least 23 different genes are associated with Fanconi anemia, including *BRCA1*, *BRCA2*, *FANCI*, *FANCD2*, *XPF*, and *SLX4*. Together, these genes constitute the Fanconi anemia DNA repair pathway, which has a critical role in repairing DNA interstrand crosslinks (ICLs) that are generated by carcinogens, metabolic by-products, or chemotherapeutic agents [69]. The Fanconi anemia pathway also protects stalled replication forks from nucleolytic degradation, thus permitting accurate DNA repair.

Interstrand crosslink repair is a highly sophisticated process that involves interplay between proteins that recognize the DNA lesion, nucleases that cleave and unhook the ICL, translesion synthesis polymerases that catalyze bypass synthesis over the unhooked ICL, and homologous recombination proteins that complete DNA repair [64]. The SLX4 scaffold has critical roles in ICL repair, stimulating XPF-ERCC1 to catalyze ICL unhooking and coordinating SLX1 and MUS81-EME1 for Holliday junction resolution in the later stages of repair [10, 11, 17, 19]. Satisfactorily, our proteomics datasets contain several Fanconi anemia proteins, namely FANCI, XPF, BRCA1, BRIP1, and BRCA2. Notably missing from our datasets is FANCD2, which forms a stable complex with FANCI. Nevertheless, we detected a plethora of proteins that function in ICL repair but are not formally designated as Fanconi anemia proteins, including ICL-processing nucleases (SNM1A), translesion synthesis polymerases (REV1, REV3L), and homologous recombination factors (e.g., MRE11, RAD50, BLM, SLX1, MUS81-EME1) [64]. The interactions and proximities between SLX4 and these different preys were detected in the absence of exogenous crosslinking agents, presumably reflecting the removal of ICLs that arise during cell growth.

### Molecular Insights into SLX4 Post-Translational Modifications

Post-translational modifications induce specific molecular changes that enable cells to respond to stimuli quickly and reversibly. Indeed, the partner proteins and subcellular localization of SLX4 are regulated by PTMs, including phosphorylation, ubiquitylation, SUMOylation, and PARylation [20, 33–36]. Consistent with these findings, we detected several enzymes that may decorate SLX4 with different PTMs. The kinase CDC7 and mitochondrial phosphatase PGAM5 appeared as high-confidence SLX4 interactors in the BioID and AP-MS datasets, suggesting that these may be relatively robust interactions. Interestingly, PGAM5 was captured in two previous AP-MS studies of SLX4 [14, 37]. Given that PGAM5 regulates mitochondrial dynamics and programmed cell death [70], it is worth exploring whether this interaction highlights non-canonical roles of SLX4 complexes in mitochondrial biology. On the other hand, CDC7 is a highly conserved serine-threonine kinase that promotes DNA replication by activating origins of replication and, thus, has a key role in promoting the G1/S phase progression [71, 72]. Whether CDC7 phosphorylates SLX4 is a question for future research.

The BioID dataset revealed several high-confidence hits between SLX4 and members of the Protein Inhibitor of Activated STAT (PIAS) family, including PIAS1, PIAS3, and PIAS4. These proteins function as E3 SUMO ligases and covalently attach SUMO to target proteins [73]. Protein SUMOylation contributes to transcriptional repression, the maintenance of heterochromatin, and DNA repair [73]. Indeed, PIAS1, PIAS3, and PIAS4 facilitate the recruitment of DNA repair proteins to DNA lesions [74, 75]. SUMO1 also appeared as a high-confidence SLX4 interactor. It is tempting to speculate that SLX4 is a substrate for PIAS family E3 ligases.

### SLX4 and Chromosome Structure: From Cohesion to the Centromere

The SLX4 interactome contains several high-confidence preys that function in chromosome structure and dynamics, most notably chromosome/chromatid cohesion. For example, we detected the vast majority of the subunits that form cohesin, a multiprotein complex containing two coiled-coil subunits, SMC1A and SMC3, the kleisin subunit RAD21, and the additional regulatory subunits SA1/2 and PDS5A/B [76, 77]. We also detected an interaction between SLX4 and NIPBL, which functions as the cohesin loading complex. The canonical role of cohesin is to ensure faithful chromosome segregation by holding sister chromatids together after DNA replication until the onset of anaphase in mitosis [76, 77]. However, cohesin also has an important role in DNA double-strand break repair, where it promotes accurate repair by HR using the sister chromatid as the template. Additionally, cohesin holds sister chromatids in close proximity at stalled replication forks to ensure a specialized type of error-free DNA repair called template-switch replication. We speculate that the high-confidence interactions between SLX4 and SMC1A, SMC3, RAD21, PDS5A/B, and NIPBL reflect their overlapping roles in DNA repair and chromosome segregation [17, 76, 77].

We also captured high-confidence interactions between SLX4 and proteins that function in the mitotic spindle, the most prominent of which are TPX2 and KIF23. The mitotic spindle is a microtubule-based assembly that separates sister chromatids during division. Centromeres are chromosomal regions that ensure that mitotic spindle fibres are correctly attached to kinetochores during cell division. Importantly, centromeres contain long stretches of repetitive alpha-satellite DNA, which are prone to DNA damage arising from secondary structures or replication-transcription conflicts [78]. Indeed, mass spectrometry analysis of Xenopus extracts reveals that centromeric DNA is enriched with DNA damage response proteins [79]. It is thus reasonable to propose that SLX4 and its associated structure-selective endonucleases resolve the branched DNA structures or recombination intermediates that arise in the centromere.

### Connections between SLX4 and Chromatin Biology

The chromatin environment is a constraint for all cellular pathways that use DNA as a substrate, including DNA replication and repair. Chromatin dynamics play a key role in maintaining genome stability and DNA repair involves substantial changes in chromatin composition and dynamics [80]. Briefly, chromatin around the DNA lesion is remodeled and relaxed by histone modifications (e.g., γH2AX) and changes in the local proteome to facilitate repair. Chromatin mobility also promotes homology search during HR [80]. The transient decompaction of chromatin is reversed when repair is complete, and the genome reverts to its original state.

The SLX4 interactome contains many proteins that function in chromatin biology. The enrichment of these factors in the BioID dataset likely reflects indirect or transient interactions with SLX4. One of the most prominent interactors in the chromatin biology cluster is YEATS2, a histone crotonylation and acetylation reader that forms part of the ADA Two A-Containing (ATAC) histone acetyltransferase complex [81–84]. YEATS2 is required for transcriptional activation of ATAC target genes [85]. This result could indicate that SLX4 is also involved in activating these target genes; additional links between SLX4 and gene transcription are discussed below.

Interestingly, over one-third of the chromatin remodeling proteins are associated with repressive chromatin environments, with notable examples being ATRX, SIRT1, and LRIF1 (for reviews, see [86, 87]). These results hint that SLX4 may have a role in chromatin compaction and heterochromatin maintenance. This is not without precedent, as previous studies established links between the Fanconi anemia pathway and chromatin state [88–90].

Satisfactorily, we captured multiple subunits from chromatin remodeling complexes, increasing our confidence in the biological relevance of these interactions. For example, BioID revealed proximal interactions between SLX4 and the ATRX-DAXX histone chaperone complex. ATRX is an ATP-dependent SWItch/Sucrose Non-Fermentable (SWI/SNF) chromatin remodeling protein that binds the histone chaperone DAXX. The ATRX-DAXX complex deposits the histone variant H3.3 onto repetitive genomic loci, including telomeres and pericentric chromatin, to maintain the repressive state [91–95]. ATRX also limits replication stress by suppressing the formation of DNA secondary structures such as G-quadruplexes [95] and maintaining the stability of common fragile sites [96]. The ATRX-DAXX complex promotes a sub-pathway of HR that involves extended DNA synthesis and crossover formation, with GEN1 and MUS81 resolving the resultant Holliday junctions [97, 98]. Given that SLX4 coordinates SLX1 and MUS81-EME1 to catalyze Holliday junction resolution [10, 11], these results imply an intriguing functional link between SLX4 and ATRX-DAXX in DNA repair.

In addition to ATRX, we identified other SWI/SNF family members as high-confidence SLX4 interactors, such as SMARCA4 (also known as BRG1). SMARCA4 is the chromatin remodeling ATPase of a larger macromolecular SWI/SNF protein complex that activates or represses the transcription of target genes [99]. Notably, the SLX4 interactome contains other components of the SMARCA4 complex, namely ACTL6A and CREBBP [100, 101]. All three subunits are high-confidence SLX4 interactors in the AP-MS datasets, implying relatively robust protein associations. CREBBP is the only chromatin remodeler captured by both techniques. CREBBP is a ubiquitously expressed transcriptional coactivator and lysine acetyltransferase with diverse roles in cell biology, including cell proliferation, DNA replication, and DNA repair [102]. As such, its association with SLX4 may reflect functions beyond the SMARCA4 complex. Collectively, the wealth of chromatin remodelers that form part of the SLX4 interactome highlight exciting new avenues for future research.

### SLX4 and RNA Biology: Transcription and Transcription Regulation

Our BioID dataset is enriched with factors that broadly function in RNA metabolism, including splicing, transcription, and transcription regulation. We envision this to reflect two emerging trends in genome stability. On the one hand, increasing evidence shows that gene transcription is a major source of endogenous replication stress and the ensuing genome instability. For example, collisions between the transcription and replication machinery can cause replication fork stalling, and highly transcribed genes are linked to increased recombination rate and mutagenesis [103]. Aberrant formation or processing of three-stranded RNA-DNA hybrids called R-loops, which frequently arise during transcription, can also compromise genome integrity [104]. It is becoming increasingly clear that DNA repair proteins, including several in the SLX4 interactome, are involved in the cellular response to these sources of genotoxic stress [103–105]. However, certain DNA repair proteins are recruited to active promoters in the absence of exogenous DNA damage, where they promote transcription [106]. For example, XPF-ERCC1 is recruited to RNA pol II promoters and is required for the initial activation of genes associated with cell growth [107, 108]. Additionally, SLX4 transcriptionally upregulates the tumor-suppressive p63 isoform and suppresses squamous cell carcinoma in mice [109]. This raises the possibility that SLX4 and its partner proteins could function similarly at other promoters. It will be important to determine how SLX4 is recruited to promoters in different cellular contexts.

In our BioID dataset, the most prominent transcriptional proteins detected in proximity to SLX4 are MGA, ZMYM4, ZNF318, CDC73 and CCNT1. MGA is a dual specificity transcription factor that recruits the non-canonical Polycomb Repressive Complex ncPRC1.6 to target promoters to repress transcription [110–112]. This provides another potential link between SLX4 and chromatin repression. There is little information about ZMYM4 and ZNF318 besides the basic characterization as transcription factors. CCNT1 forms part of the P-TEFb (Positive Transcription Elongation Factor), which positively regulates transcriptional elongation and is required for the transcription of most class II genes [113]. Notably, we also detected high-confidence interactions between SLX4 and several components of the Polymerase-Associated Factor 1 Complex (PAF1C), representing another transcription elongation factor of RNA pol II. Mammalian PAF1C contains six core subunits, namely PAF1, CDC73, CTR9, LEO1, RTF1, and SKI8, and has diverse functions that positively regulate gene expression genome-wide [114, 115]. Of these six subunits, PAF1 and CDC73 were detected as high confidence SLX4 interactors by BioID and AP-MS, while AP-MS uniquely captured CTR9. It is tempting to speculate that SLX4 and its nuclease partners facilitate transcription elongation by resolving replication-transcription conflicts. Overall, the SLX4 interactome contains a diverse array of proteins that function in RNA metabolism, stressing the connection between gene expression and genome stability.

In summary, we leveraged complementary techniques, BioID and AP-MS, to establish the first comprehensive SLX4 interactome. This network contains candidate proteins that could interact with SLX4 directly, indirectly (mediated by another protein), or spatially (within the radius of biotinylation). These partner proteins may regulate the known functions of SLX4 in nuclear DNA repair or confer novel functions in other cellular processes. Unlike previous AP-MS methods that only characterized stable SLX4-complexes, we increased the exploration space by using BioID in parallel to capture weak and transient interactions within the living cell. In doing so, our datasets provide a wealth of insight into the cellular environment and functions of SLX4. The candidate proteins collectively allow for a deeper understanding of how SLX4 functions in DNA repair while revealing new cellular processes that may involve SLX4. Our findings will pave the way for future research to explore understudied relationships between SLX4 and RNA metabolism, chromosome structure and dynamics, and chromatin biology.

## ACKNOWLEDGMENTS

We thank members of the Wyatt and Gingras labs for their helpful suggestions and advice throughout this project. We also thank members of the PhD thesis committee for C.A., including Grant Brown and Eric Campos. The authors wish to thank Kimberly Lau at the SickKids Imaging Facility (The Hospital for Sick Children, Toronto, Canada) for excellent technical assistance and experimental support. This work was supported by a Natural Sciences and Engineering Research Council of Canada (NSERC) Discovery Grant (RGPIN-2017-06670 to H.D.M.W. and RGPIN-2019-06297 to A.-C.G.). Proteomics was performed at the Network Biology Collaborative Centre at the Lunenfeld-Tanenbaum Research Institute, a facility supported by Canada Foundation for Innovation funding, by the Ontarian Government, and by Genome Canada and Ontario Genomics (OGI-139). C.A. holds an Ontario Graduate Scholarship and is a previous recipient of the Frederick Banting and Charles Best Canada Graduate Scholarship (Master’s). B.J.A.D. was supported by an NSERC Postgraduate Scholarship (Doctoral). H.D.M.W. holds a Tier II Canada Research Chair in Mechanisms of Genome Stability (#950-231487). We apologize to our colleagues whose work we were unable to cite due to space limitations.

## AUTHOR CONTRIBUTIONS

Conceptualization, C.M.A. and H.D.M.W.; Methodology, C.M.A., B.J.A.D., V.H.W.C., C.J.W., M.P., A.-C.G., and H.D.M.W.; Software, C.M.A., B.J.A.D., C.J.W., and A.-C.G.; Investigation, C.M.A., B.J.A.D., V.H.W.C., C.J.W., M.P., and H.D.M.W.; Writing – Original Draft, C.M.A. and H.D.M.W.; Writing – Review & Editing, C.M.A., B.J.A.D., V.H.W.C., C.J.W., M.P., A.-C.G., and H.D.M.W.; Visualization, C.M.A. and H.D.M.W.; Supervision, A.-C.G. and H.D.M.W.; Funding Acquisition, A.-C.G. and H.D.M.W.

## SUPPLEMENTAL FIGURE LEGENDS

**Figure S1.**
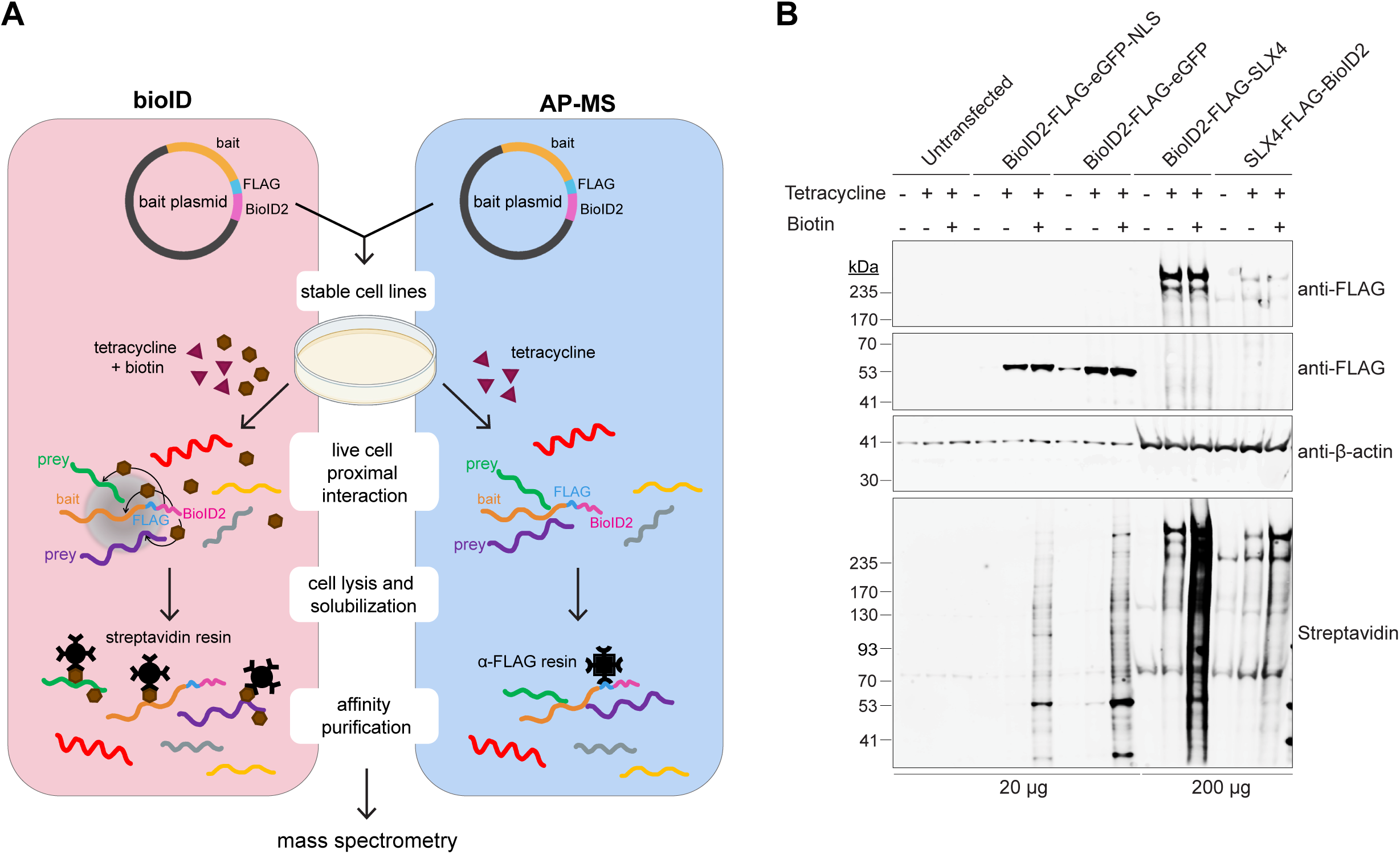
Validating the expression and functionality of BioID2-FLAG constructs for proximity-dependent labeling and affinity purification mass spectrometry. **(A)** Workflow for identification of protein partners using proximity-dependent biotinylation (BioID) and affinity purification coupled to mass spectrometry (AP-MS). Bait plasmids contain the gene-of-interest with an N- or C-terminal BioID2-FLAG tag and are stably integrated into Flp-In T-REx HEK293 cells for conditional expression with tetracycline (red diamonds). In BioID, cell cultures are supplemented with exogenous biotin (brown pentagons) for live cell labelling. After optimized cell lysis, bait proteins are enriched on streptavidin (BioID) or anti-FLAG (AP-MS) resin. Prey proteins are identified by mass spectrometry. **(B)** Western blot analysis of whole cell extracts from Flp-In T-REx HEK293 cells containing the integrated constructs. Untransfected cells are shown for comparison. Cell cultures were left untreated, incubated with tetracycline for 25 hr to induce expression of the integrated construct, or incubated with tetracycline for 25 hr and biotin (50 μM) for 8 hr to induce construct expression and live cell biotinylation of proximal proteins, respectively. The BioID2-FLAG-eGFP and BioID2-FLAG-eGFP-NLS constructs were induced with 0.001 μg/mL tetracycline, whereas BioID2-FLAG-SLX4 and SLX4-FLAG-BioID2 were induced with 1 μg/mL tetracycline. The indicated amount of cell extract was analyzed by western blotting with α-FLAG and IR800-conjugated streptavidin antibodies; β-actin was used as a loading control.

**Figure S2.**
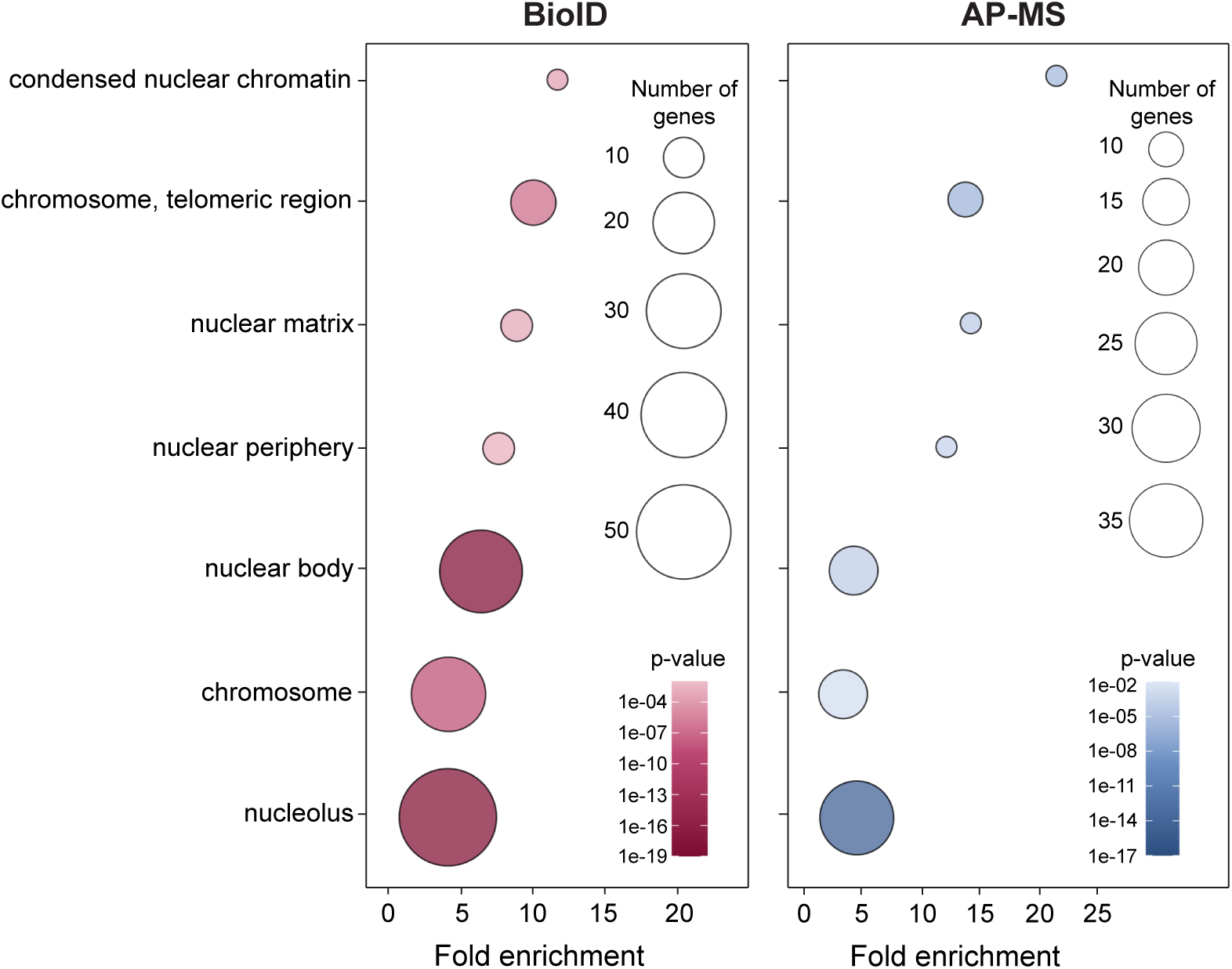
Cellular component gene ontology (GO) analysis of high-confidence SLX4 interactors identified by BioID and AP-MS. Bubble plot displaying cellular component GO terms associated with SLX4 binding proteins captured by BioID (red) or AP-MS (blue). Terms were selected from Revigo after applying a medium list cut-off (0.7) [58]. P-values as listed by Princeton Generic GO Term finder [57]. Fold enrichment represents the number of genes associated with a specific GO function in each proteomics dataset compared to the number of genes related to that function in the human genome.

